# PDS5 proteins control genome architecture by limiting the lifetime of cohesin-NIPBL complexes

**DOI:** 10.1101/2025.08.30.673243

**Authors:** Gordana Wutz, Iain F Davidson, Edward J Banigan, Ryotaro Kawasumi, Roman R Stocsits, Wen Tang, Kota Nagasaka, Lorenzo Costantino, Ralf Jansen, Kouji Hirota, Dana Branzei, Leonid A Mirny, Jan-Michael Peters

**Affiliations:** Research Institute of Molecular Pathology (IMP), Vienna Biocentre (VBC), Campus-Vienna-Biocenter 1, 1030 Vienna, Austria; Institute for Medical Engineering and Science, Department of Physics, Massachusetts Institute of Technology, Cambridge, MA, USA; The AIRC Institute of Molecular Oncology Foundation, IFOM ETS, Via Adamello 16, Milan 20139, Italy; Department of Chemistry, Graduate School of Science, Tokyo Metropolitan University, 1-1 Minamiosawa, Hachioji-shi, Tokyo 192-0397, Japan; Institute of Molecular Biotechnology of the Austrian Academy of Sciences (IMBA), Vienna BioCenter (VBC), Austria; University of Duisburg-Essen, Research Center One Health Ruhr, University Alliance Ruhr, Universität Straße 2, 45141, Essen Germany

**Keywords:** compartmentalization, CTCF boundaries, genome organization, loop extrusion, polymer simulations, single-molecule imaging, WAPL

## Abstract

Cohesin-NIPBL complexes extrude genomic DNA into loops that are constrained by CTCF boundaries. This process has important regulatory functions and weakens the separation between euchromatic and heterochromatic compartments. Cohesin can also bind PDS5A or PDS5B, which do not support loop extrusion but are required for the formation of CTCF boundaries. How PDS5 proteins perform this function is unknown. Here we show by *in vitro* single-molecule imaging that PDS5 proteins stop loop extrusion by facilitating the dissociation of NIPBL from cohesin. Hi-C experiments suggest that this function is required for the establishment of CTCF boundaries in cells. *In silico* modelling indicates that PDS5 proteins enable the separation between compartments by limiting cohesin’s velocity and chromatin-residence time. The degree of this compartmentalization depends on the frequency with which chromatin is extruded relative to the time it takes for compartments to form. These results identify PDS5 proteins as key regulators of genome organization.

**Highlights:**

- PDS5 proteins stop loop extrusion by facilitating dissociation of NIPBL from cohesin.
- PDS5 proteins strengthen CTCF boundaries by limiting the lifetime of cohesin-NIPBL.
- PDS5 proteins regulate compartmentalization.
- Compartmentalization is governed by polymer relaxation and loop extrusion dynamics.

## Introduction

Cohesin complexes extrude chromatin fibers into loops, which have important functions in genome regulation^1^. The loops formed by cohesin (“cohesin loops”) promote the assembly of antigen receptor genes by V(D)J recombination^2^, DNA repair by homologous recombination^3, 4^, the spreading of histone modifications and histone modifying enzymes^4–6^, the local separation of sister chromatids^7^ and contribute to the timing of DNA replication^8, 9^. Although cohesin is dispensable for short-range enhancer-promoter interactions^10–12^, cohesin loops are essential for the activation of selected genes during development, cell differentiation and stimulation^13–17^. Defects in these processes are thought to be the cause of “cohesinopathies”^18, 19^ and contribute to tumorigenesis^20, 21^. Furthermore, cohesin loops suppress the segregation of chromatin into A and B compartments^22–27^, megabase-sized domains that correspond to transcriptionally active euchromatic regions and inactive heterochromatic regions, respectively^28^. Cohesin loops could therefore be important for disrupting spurious local chromatin structures^29^.

Key for many of these functions are regulatory mechanisms that control cohesin-mediated loop extrusion in space and time. Some of these mechanisms have been identified, but others remain to be understood. The cohesin release factor WAPL limits the residence time of cohesin on chromatin^30, 31^ and thus determines the length and lifetime of cohesin loops^22, 23, 26^. In developing B cells and neurons, WAPL is transcriptionally downregulated to enable the generation of exceptionally long cohesin loops, which promote V(D)J recombination and protocadherin promoter choice, respectively^16, 32, 33^.

Spatially, the positions of cohesin loops in the genome are controlled by boundary elements that halt loop extrusion. Cohesin complexes can presumably associate with DNA anywhere in the genome, with some preference for loading at enhancers^34–36^, but relocalize through extrusion and then accumulate at boundary elements. The most prominent type of these boundaries is formed by the DNA binding protein CTCF^24–26, 37–41^, but active genes, the replicative helicase MCM, R-loops and cohesin complexes that mediate cohesion between replicated sister chromatids are also barriers for loop-extruding cohesin^42–47^. CTCF functions as an asymmetric boundary that positions cohesin loops between convergent CTCF binding sites^26, 44, 47–51^. In Hi-C studies of large cell populations, these CTCF sites appear as boundaries between topologically associating domains (TADs)^52, 53^ and as “anchors” for corner peaks^54^. Both TADs and corner peaks are emergent features of cell population studies, i.e. are patterns that are formed by the superimposition of cohesin loops in individual cells^55–57^. CTCF sites are essential for the functions of cohesin loops in gene regulation and V(D)J recombination^58–60^ and can be inactivated by DNA methylation. For example, methylation of CTCF binding sites in germ cells establishes imprinted gene expression patterns during mammalian development^61^ and demethylation of CTCF binding sites determines protocadherin promoter choice in developing neurons^62^.

Cohesin’s loop extrusion activity depends on NIPBL^25, 63, 64^ (called Scc2 in yeast). NIPBL is a 316 kDa HEAT repeat protein that forms hetero-dimers with MAU2, promotes cohesin’s association with DNA^65, 66^ and stimulates cohesin’s ATPase activity, which is essential for loop extrusion^63, 64, 67, 68^. Interactions between NIPBL-MAU2 and cohesin are relatively short-lived (∼1 min)^63, 69, 70^ and depend on a binding site on cohesin’s kleisin subunit SCC1 (also called RAD21 or Mcd1)^71^.The same site on SCC1 can be bound by PDS5 proteins^71^, HEAT repeat proteins that are structurally similar to NIPBL but are unable to stimulate cohesin’s ATPase and loop extrusion activities^63, 68^. Vertebrate genomes encode two PDS5 paralogs, PDS5A and PDS5B^72^. Yeast Pds5 can inhibit the ability of NIPBL/Scc2 to stimulate cohesin’s ATPase activity, suggesting that NIPBL and PDS5 proteins compete for binding to cohesin^68^.

PDS5 proteins are essential for mouse development^73^ and are required for sister chromatid cohesion, replication fork progression and protection, assembly of mitotic and meiotic chromosomes and meiotic recombination^74–77^. At the molecular level, several functions of PDS5 proteins have been identified. PDS5 proteins bind to WAPL and contribute to the release of cohesin from chromatin^30, 78–82^, but can also promote loading of cohesin onto DNA *in vitro*^83^. PDS5 proteins further facilitate acetylation of cohesin’s SMC3 subunit by the acetyltransferases ESCO1 and ESCO2^80, 84, 85^. SMC3 acetylation is required for sister chromatid cohesion mediated by cohesive cohesin complexes^86, 87^. Acetylation can also occur on loop-extruding cohesin complexes^88, 89^, which reduces their ability to form long chromatin loops in a manner that depends on PDS5^88, 90, 91^. It has therefore been proposed that PDS5A functions as a “brake” for loop-extruding cohesin^91^. In yeast, the only essential function of Pds5 is to recruit the SUMO protease Ulp2 to de-sumoylate cohesin, which otherwise would be targeted for degradation by SUMO chains^92^.

Unexpectedly, we observed previously that depletion of PDS5 proteins reduces TAD boundaries and corner peaks^26^ suggesting that these proteins are also required for the formation or function of CTCF boundaries. Here, we have used genomic approaches in human and chicken cells, biochemical reconstitution, single-molecule imaging *in vitro*, and *in silico* modelling to understand how PDS5 proteins perform this function. Our results indicate that PDS5 proteins stop loop extrusion by promoting the dissociation of NIPBL from cohesin and that this function is required for the establishment of efficient CTCF boundaries. Simulations further suggest that PDS5 proteins control the average speed of extrusion by cohesin, which accelerates when PDS5 proteins are depleted. The simulations also suggest that PDS5 proteins and WAPL modulate compartmentalization by affecting both loop extrusion and chromatin fiber conformational dynamics, therefore shifting the competitive balance between these two mechanisms. These results indicate that PDS5 proteins control key features of genome organization by limiting the lifetime of cohesin-NIPBL complexes, and the average speed of extrusion.

## Results

### PDS5 proteins are required for CTCF boundaries

We had previously observed that co-depletion of PDS5A and PDS5B by RNA interference (RNAi) reduced TAD insulation and the number of corner peaks^26^ To study the mechanistic basis of these phenotypes and to rule out off-target effects, we generated HeLa cell lines in which PDS5A and PDS5B could be depleted singly or in combination within five hours by auxin-induced degradation (Figures S1A-S1D). Hi-C analysis of these cells synchronized in G1 phase revealed that depletion of PDS5A and PDS5B alone reduced the number of detectable corner peaks but had only minor effects on the length of chromosomal *cis*-interactions (Figures S1E-S1I; identical numbers of sequence reads were used for comparisons in this and all other Hi-C experiments; Table S1), as reported^93^. In contrast, simultaneous depletion of both proteins (hereafter called PDS5A/B) greatly increased the length of chromosomal *cis*-interactions and reduced the TAD insulation score and corner peak number more strongly (Figures 1A-D and S1I), consistent with the notion that PDS5A and PDS5B have partially redundant functions^73^. The reduction of corner peaks and TAD insulation in PDS5A/B depleted cells supports the hypothesis that PDS5 proteins are required for the formation or function of CTCF boundaries^26^.

**Figure 1.**
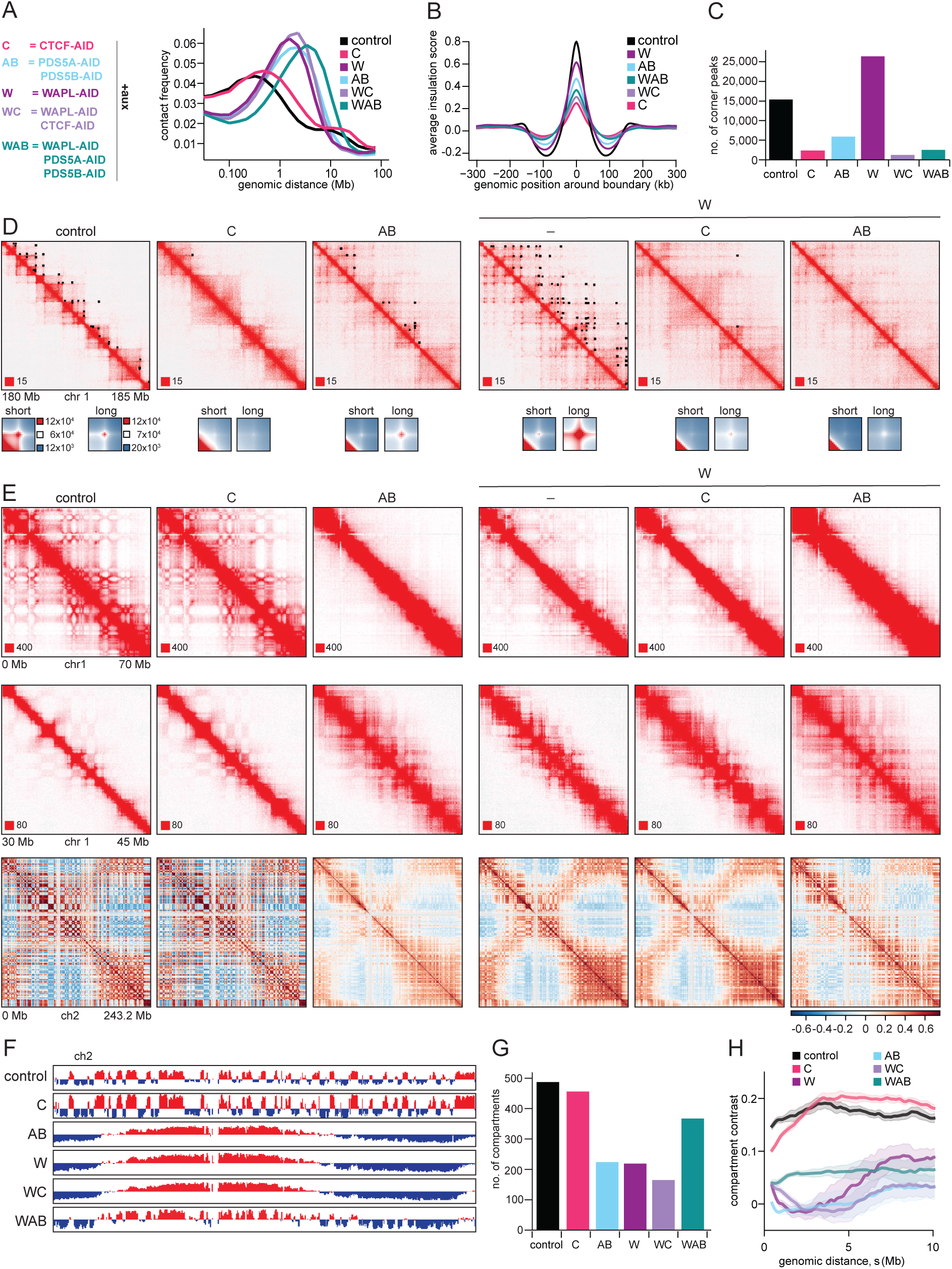
PDS5 proteins are essential for maintaining CTCF boundary integrity and proper genome compartmentalization. **A.** *Left:* Abbreviations of AID-tagged cell lines treated with auxin in this study. *Right:* Intra-chromosomal contact frequency distribution as a function of genomic distance for control cells and cells depleted of, CTCF (C), WAPL (W), PDS5A/B (AB), WAPL/CTCF (WC) and WAPL/PDS5A/PDS5B (WAB). **B.** Average insulation score around TAD boundaries identified in control cells for the same conditions as in (A). **C.** Number of corner peaks identified by HICCUPS for the same conditions as in (A). **D.** *Top:* Coverage-corrected Hi-C contact matrices for the 180–185 Mb region of chromosome 1, visualized using Juicebox under the same conditions as in (A). Corner peaks identified by HICCUPS are marked with black rectangles in the upper triangle. *Bottom:* Total contact counts around corner peaks are shown for different conditions: corner peaks ≤350 kb (short) identified in control and corner peaks >350 kb (long) identified in WAPL-depleted cells. Matrices are centered at corresponding loop anchor sites in vertical and horizontal orientation. **E.** *Top:* Coverage-corrected Hi-C contact matrices for the 0–70 Mb region of chromosome 1, visualized using Juicebox under the same conditions as in (A). Notably, the checkerboard pattern at long distances is significantly diminished following WAPL and WAPL/CTCF depletion and completely absent after PDS5A/B and WAPL/PDS5A/PDS5B (WAB) depletion. Middle: Coverage-adjusted Hi-C contact matrices for the 30–45 Mb region of chromosome 1, visualized using Juicebox under the same conditions as in (A). The strongest checkerboard pattern, which is at this resolution close to the diagonal, appears following WAPL/PDS5A/PDS5B (WAB) depletion. *Bottom:* Pearson correlation matrix for the whole chromosome 2. **F.** Compartment contrast as a function of genomic distance (s) for control and perturbation conditions (see Fig. S2G and Methods). Shaded areas indicate standard error of the mean, calculated from the distance-dependent compartment contrasts of each individual chromosome. **G.** The corresponding compartment signal tracks at 500 kb bin resolution are shown. The tracks were plotted using Juicebox. **H.** Genome-wide quantification of compartmental switches.

To understand how PDS5 proteins perform these functions, we generated cell lines in which CTCF or WAPL could be depleted by auxin-induced degradation (Figures S1A-S1D and S1J-S1L) and compared the Hi-C phenotype of PDS5A/B depleted cells with those depleted of CTCF or WAPL. These comparisons revealed that PDS5A/B depleted cells display distinct subsets of the phenotypes observed in CTCF and WAPL depleted cells^23, 26, 37, 94^. PDS5A/B depleted cells resembled CTCF depleted cells insofar as corner peaks were reduced in both (Figures 1C and 1D), whereas they resembled WAPL depleted cells with respect to the length of chromosomal *cis*-interactions, which were greatly increased in both PDS5A/B and in WAPL depleted cells (Figures 1A and S1M). In contrast, *cis*-interactions were lengthened to only a small extent in the absence of CTCF (Figures 1A and S1M), and corner peak numbers did not decrease but instead *increased* in the absence of WAPL (Figures 1C and 1D). The observation that PDS5A/B depleted cells share distinct features with both CTCF depleted cells (reduced corner peak numbers) and WAPL depleted cells (elongated *cis*-interactions) suggests that PDS5 proteins control two different properties of cohesin-mediated loop extrusion: (i) the stalling of loop-extruding cohesin at CTCF sites, a process that is responsible for the formation of corner peaks, and (ii) cohesin’s residence time on chromatin, a property that determines the processivity of loop extrusion.

To further characterize the roles of PDS5 proteins in corner peak formation, we generated cell lines in which WAPL could simultaneously be co-depleted either with PDS5A/B or with CTCF (Figures S1A-S1D). These experiments revealed that the strong increase in corner peak numbers caused by WAPL depletion was fully prevented by co-depletion of PDS5 proteins (Figures 1C and 1D). In this respect, PDS5A/B depletion resembled the effect of CTCF depletion, which also prevented the formation of corner peaks in WAPL depleted cells (Figures 1C and 1D). The similarity of these effects supports the notion that not only CTCF but also PDS5 proteins are required for CTCF boundaries.

While the effects of PDS5A/B depletion on corner peaks were epistatic to those of WAPL depletion, the effects of PDS5A/B depletion on the length of chromosomal *cis*-interactions were additive with those of WAPL depletion. Triply depleted cells contained even longer chromatin loops (average size 645 kb) than cells depleted of only WAPL (412 kb) or PDS5A/B (378 kb), compared to an average loop size of 119 kb in control cells (Figure S1M; average loop sizes were estimated from the derivative of the genome-wide contact probability curve, P(s), in log-log space, i.e. the slope of log(P(s))^22, 95, 96^. This result suggests that co-depletion of WAPL and PDS5A/B increases the chromatin-residence time and/or velocity of loop-extruding cohesin more than depletion of WAPL or PDS5 proteins alone (see below).

### PDS5 proteins control chromatin compartments

Depletion of cohesin or NIPBL leads to stronger and more segmented compartmentalization^24–, 26^, whereas depletion of WAPL or PDS5 proteins weakens compartmentalization^22, 23, 26^, indicating that loop extrusion suppresses compartmentalization^27^.

We therefore assumed that co-depletion of PDS5A/B and WAPL would reduce compartments even further. Unexpectedly, however, depletion of all three proteins had differential effects on long-range compartments (>5 Mb Figure 1E, top panels) and short-range compartments (<5 Mb; Figure 1E, middle panels). In cells depleted of PDS5 proteins or WAPL alone, the checkerboard pattern of long-range compartments was reduced in Hi-C maps (Figure 1E, top panels) and Pearson correlation maps (Figure 1E, bottom panels). Eigenvector analyses indicated that A and B compartments became larger under these conditions (Figure 1F). In contrast, in cells co-depleted of WAPL and PDS5A/B, short-range compartmental contacts were partially recovered compared to cells depleted of WAPL or PDS5 proteins alone (Figure 1E, middle panels). These checkerboard patterns correlated with alternating patterns of euchromatic and heterochromatic histone modifications (H3K4me1 and H3K9me3, respectively; Figure S1N), indicating that they represent *bona fide* A and B compartments. Eigenvector analysis showed that these A and B compartments had approximately the same lengths as in control cells (Figure 1F). Correspondingly, the number of A and B compartments in cells co-depleted of WAPL and PDS5A/B (367) was closer to the number in control cells (487) than to those in cells depleted of WAPL or PDS5A/B alone (219 and 225, respectively; Figure 1G).

We quantified compartmentalization by measuring the distance-dependent compartment contrast^27^. This metric calculates the relative intensity of intra-compartmental contacts (AA and BB) compared to inter-compartmental contacts (AB), and it does so separately at each genomic distance (i.e., along each Hi-C diagonal; see Methods). As observed in the Hi-C maps, depletion of WAPL and PDS5 proteins individually suppressed compartmentalization compared to control conditions across all genomic distances, particularly at length scales of a few Mb, whereas co-depletion of all three proteins resulted in a partial recovery of compartmentalization of loci separated by less than ∼5 Mb (Figure 1H and S2G). Co-depletion of PDS5 proteins and WAPL therefore prevents the suppression of short-range compartments that is otherwise seen when PDS5A/B or WAPL are depleted alone.

### Compartmentalization is controlled by cohesin’s residence time and loop extrusion velocity

To explain this phenomenon, we studied an *in silico* polymer model that recapitulates key features of genome organization^25, 27^. The model simulates a 60 Mb chromosome containing three repeats of a 20 Mb region of human chromosome 1, patterned with A/B compartments and folded by loop extrusion (Figure 2A). The model assumes that compartments are formed by affinity interactions between B-type polymer subunits, leading to the separation of A and B regions^27, 97^. Loop-extruding cohesin complexes randomly bind to the polymer fiber and progressively enlarge loops until they either stall at barriers (such as other cohesin complexes) or stochastically unbind^98, 99^. In simulations, when cohesin’s chromatin residence time is long, cohesin accumulates in vermicelli (Figure 2A, bottom), as experimentally observed in PDS5A/B-depleted cells^26^ (Figure S2E). To investigate the effects of loop extrusion on compartmentalization, we performed polymer simulations with different cohesin residence times (*t*_res_), extrusion velocities (*v*), and mean distances between cohesin complexes (*d*=1/(linear density)).

**Figure 2.**
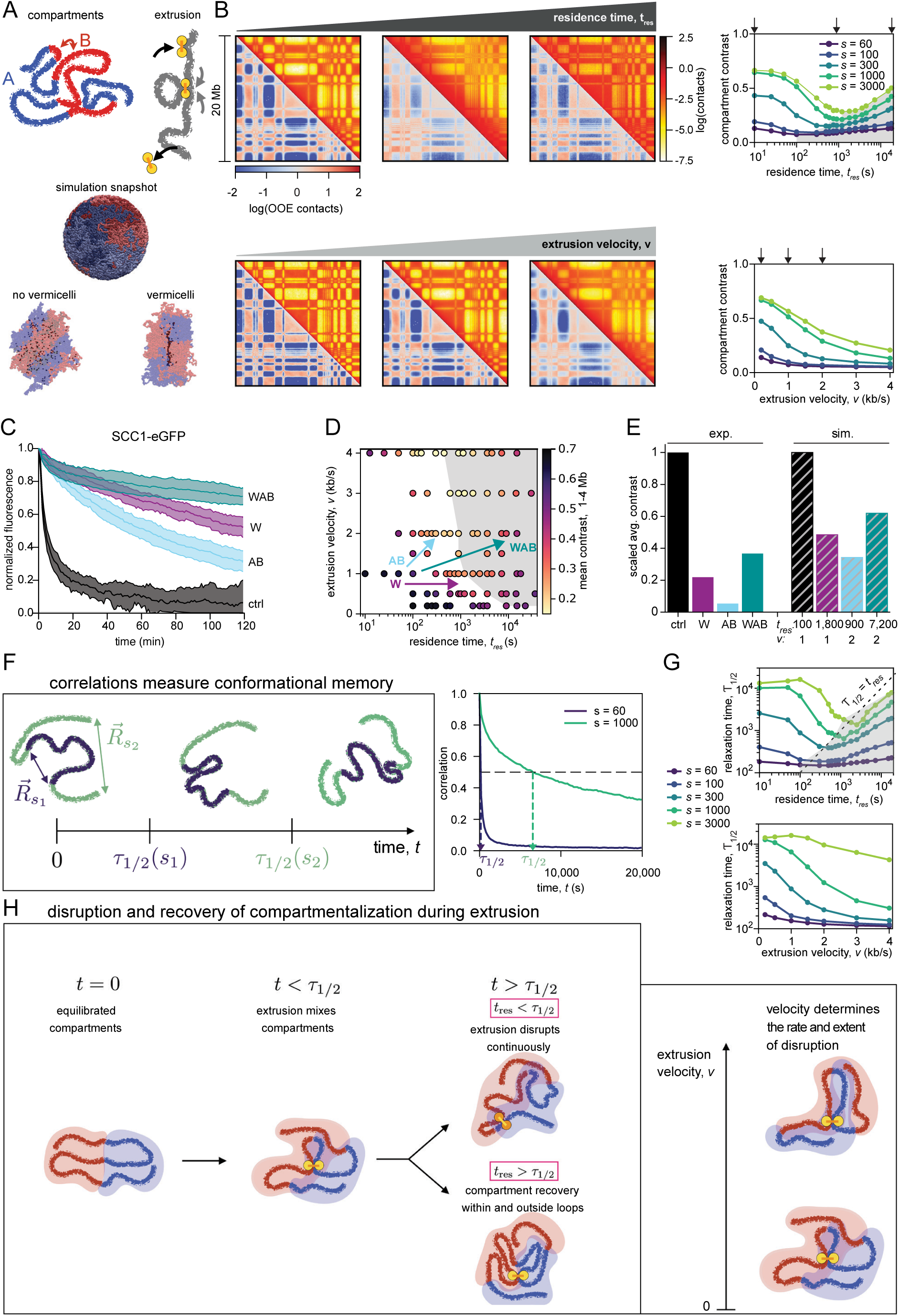
Compartmentalization is governed by the relationship between polymer relaxation and loop extrusion dynamics. **A.** Illustration of polymer simulation model and a simulation snapshots. *Top left*: The chromatin fiber compartmentalizes via affinity between monomers in B-type segments (red), which aggregates B regions while expelling neutral A regions (blue). Top *right:* Loop extruders representing cohesin (yellow) randomly bind to the chromatin fiber (gray), progressively extrude chromatin loops at speed *v*, and stochastically unbind after residence time *t_res_*. *Middle:* A simulation snapshot shows microphase-separated A and B regions of a chromosome in a spherical volume (blue and red, respectively), with the color darkening from one end of the chromosome to the other such that comparable darkness of color indicates genomic proximity. *Bottom:* Simulation snapshots from example simulations performed without spherical confinement (in periodic boundary conditions). Transparent blue and red indicate A and B compartments, respectively, while solid black subunits are cohesins. Left shows a chromosome simulated with cohesin residence time *t_res_*=100 s, which does not exhibit the vermicelli phenotype. Right shows a simulation with *t_res_*=7200 s, displaying vermicelli. **B.** *Left:* Contact maps from simulations with different extrudercohesin residence times, *t_res_* (top row), and extrusion velocities, *v* (bottom row), with *t_res_* or *v* increasing from left to right. For each row, only a single extrusion variable (*t_res_* or *v*) is varied. Lower left half of each map shows observed-over-expected contacts, while upper right shows unnormalized contact frequency. *Right:* Compartment contrast versus *t_res_* (top) and *v* (bottom) for contacts between sites at different genomic distances, *s* (given in kb and indicated by different colors), along the chromatin polymer. Arrows denote parameters used for simulation maps to the left. The gray region marks the regime in which compartment contrast increases with increasing residence time. For all simulations *d*=500 kb, *t_res_*=100 s except where *t_res_* is varied, and *v*=1 kb/s except where *v* is varied. **C.** Normalized fluorescence signal of GFP-tagged SCC1 over time following photobleaching, measured in iFRAP experiments in control cells and cells depleted of PDS5A/B (AB), WAPL (W), or both WAPL and PDS5A/B (WAB). Data are presented as mean ± SD; *n* > 20 cells per condition. **D.** Phase diagram of mean compartment contrast across the *t_res_*-*v* parameter space. The gray region corresponds to the area where compartment contrast increases with increasing *t_res_*. Arrows indicate predicted qualitative changes from control induced by WAPL (W), PDS5A/B (AB), and triple depletion (WAB). **E.** Bar graph showing mean compartment contrast over 1-4 Mb genomic distances in experiments and representative simulations. **F.** *Left:* Illustrations showing the physical meaning of the relaxation time, τ_1/2_(s). The sequence shows the typical time evolution of a chromosomal polymer segment, with a short segment of genomic length, *s1*, and a long segment of genomic length, *s2* (blue and green, respectively). Corresponding segment end-to-end vectors, 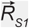 and 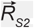, at t=0 are indicated. After time τ_1/2_(*s1*), the shorter (blue) segment has relaxed (rearranged), while the longer (green) segment retains memory of its original conformation. After time τ_1/2_(*s2*), the longer polymer segment has also lost the memory of its original conformation. *Right:* Polymer segment correlations, 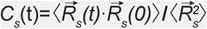 for a short (60 kb, purple) and long (1000 kb, green) polymer segments, computed from simulations with tres=100 s and v=1 kb/s. The relaxation time, τ_1/2_(s), for genomic length s is given by the time at which the corresponding correlation curve crosses 0.5. **G.** Relaxation times, τ_1/2_(s), versus residence time, tres (top), and extrusion velocity, v (bottom) for different genomic lengths, s (different colors). Gray region is the same as gray region in which compartment contrast increases in (B). **H.** Illustration depicting how chromatin compartmentalization depends on polymer relaxation time, τ_1/2_, extrudercohesin residence time, tres, and extrusion velocity, v.

Surprisingly, we found that compartment contrast, measured as in the experimental Hi-C data (Figure 1H, S2G), has a non-monotonic relationship with residence time (Figure 2B, top row). Initially, compartmentalization across all genomic distances, *s*, decreases as cohesin’s residence time is increased, consistent with previous WAPL depletion experiments and simulations^22, 23, 26, 27^ (Figure 2B). However, when residence times are increased further, compartmentalization strengthens, especially at megabase scales (Figures 2B, grey region in top right panel, and S2A). Remarkably, this strengthening of compartmentalization (Figure 2B) can occur at length scales even greater than the sizes of extruded loops (Figure S2B).

This recovery of compartmentalization at long residence times resembles the experimental effect of WAPL-PDS5A/B co-depletion. This result therefore suggests that co-depletion of WAPL and PDS5 proteins enables the formation of short-range compartments by increasing cohesin’s chromatin residence time. Consistent with this possibility, previous experiments in which WAPL and PDS5A/B had been depleted by RNAi had indicated that co-depletion of all three proteins increases cohesin’s residence time more than depletion of either WAPL or PDS5A/B alone^26^. However, this result had been unexpected since WAPL and PDS5 proteins are thought to form hetero-dimers that release cohesin from chromatin through the same pathway^30, 79, 80^. Depletion of WAPL, PDS5 proteins, or all three would thus be expected to have similar and not additive effects on cohesin’s residence time, if depletion efficiencies are comparable. We therefore performed new measurements of cohesin’s residence time by inverse fluorescence-recovery-after-photobleaching (iFRAP) in cells in which WAPL and PDS5A/B were depleted by auxin-induced degradation. Under these conditions, co-depletion of WAPL and PDS5A/B increased cohesin’s residence time much more (*t*_res_ = 12.2 hrs) than depletion of WAPL (*t*_res_ = 4.0 hrs) or PDS5A/B alone (*t*_res_ = 2.2 hrs; Figures 2C, S2C and S2D). Correspondingly, cohesin accumulated more strongly in vermicelli following co-depletion of WAPL and PDS5A/B than after depletion of WAPL or PDS5A/B alone (Figures S2E and S2F). These results support the hypothesis that co-depletion of WAPL and PDS5 proteins enables the formation of short-range compartments because cohesin has an exceptionally long chromatin residence time under these conditions. These findings further indicate that in cells co-depleted of PDS5A/B and WAPL, cohesin can only be released very infrequently from DNA, and that extruded loops must therefore be long-lived and largely static. Finally, these results raise the unexpected possibility that WAPL and PDS5 proteins have partially independent roles in releasing cohesin from chromatin (see Discussion), consistent with the observation that WAPL and PDS5 proteins can bind cohesin independently of each other^100, 101^.

Although the exceptionally long cohesin residence time in cells co-depleted of WAPL and PDS5A/B correlates well with the presence of short-range compartments, suppression of compartmentalization by either PDS5A/B depletion or WAPL depletion cannot be explained by changes in cohesin residence time alone. WAPL depletion increases cohesin’s residence time more (*t*_res_ = 4.0 hrs) than PDS5A/B depletion (*t*_res_ = 2.2 hrs), yet PDS5A/B depletion suppresses compartmentalization more than WAPL depletion (Figures 1E and H). We therefore investigated whether other properties of loop extrusion could explain the experimental observations by performing simulations with different extrusion velocities, *v*, or mean distances, *d*, between cohesin complexes. Unlike changes in residence time, increases in velocity and decreases in mean distances have monotonically suppressive effects on compartmentalization (Figures 2B, bottom panels, and S2A, S2H and S2I). In the depletion experiments, we expect that the mean distance between cohesin complexes on chromatin decreases due to the increase in cohesin’s residence time, resulting in a larger number of cohesin complexes on chromatin. The mean distance between cohesin complexes, *d*, should thus decrease more after WAPL depletion than after PDS5A/B depletion. Therefore, changes in mean distance between cohesin complexes cannot account for the larger suppression of compartmentalization by PDS5A/B depletion than by WAPL depletion. We therefore tested whether depletion of PDS5 proteins might suppress compartmentalization by increasing extrusion velocity.

To explore this possibility, and to test whether changes in extrusion velocity would be consistent with the results of our WAPL and PDS5A/B co-depletion experiments, we constructed a phase diagram for Mb-scale compartmentalization. For this purpose, we averaged compartment contrast in the 1-4 Mb range in simulations and plotted it as a function of cohesin residence times, *t*_res_, and extrusion velocities, *v* (Figure 2D). We found that increasing residence time and velocity together can reduce compartmentalization as much as a larger increase in residence time alone. These results indicate that depletion of PDS5A/B and WAPL can suppress compartmentalization to a similar degree, but by impacting different aspects of extrusion. PDS5A/B depletion could suppress compartmentalization by increasing the velocity and the residence time. WAPL depletion achieves the same effect by increasing the residence time by a larger amount, while leaving velocity unchanged (Figure 2E). Our simulations therefore suggest that PDS5 proteins might control the average speed of loop extrusion (see below for experimental tests of this hypothesis). Furthermore, we observed that extreme increases in residence time could explain the perplexing recovery of compartmentalization that we observed with WAPL and PDS5A/B co-depletion (gray region in Figure 2D). This analysis suggests that co-depletion of WAPL and PDS5A/B could enable compartmentalization because the increase in residence time is so large that it can overcome the suppressive effect of increased extrusion velocity (Figure 2E).

### Compartmentalization is governed by the relationship between polymer relaxation and loop extrusion dynamics

These results indicate that compartmentalization is suppressed by increased loop extrusion velocity and depends non-monotonically on cohesin’s residence time, with the maximal suppression for intermediate residence times. However, the origins of these effects, and more generally, the biophysical mechanisms through which loop extrusion alters compartmentalization are poorly understood. It has previously been proposed that extending cohesin-mediated loops by increasing either *t*_res_ or *v* brings together chromosomal segments from different compartments, thus weakening compartmentalization^25, 27^. However, several lines of evidence indicate that extrusion affects compartmentalization not only by bringing together distal loci, but also by altering polymer dynamics.

The formation of exceptionally long cohesin loops in cells co-depleted of WAPL and PDS5A/B does not fully suppress compartmentalization (Figure 1E and 1H), suggesting that extrusion influences compartmentalization through a mechanism other than imposing new contacts via looping. This view is supported by the surprising non-monotonic dependence of compartmentalization on cohesin residence time in our simulations (Figure 2B, top panels): while the mean loop size monotonically increases (Figure S2B), compartmentalization first decreases and then rises with increasing residence time. Furthermore, we observed that compartmentalization is not solely dependent on the static loop architecture: in simulations with loops formed at a specific residence time and subsequently frozen in place (see Methods), compartmentalization is not strongly perturbed by increasing loop sizes (Figure S2K). In contrast, maintaining the same loop architecture (i.e. constant *d* and processivity, *λ*=2*vt*res) while increasing extrusion velocity suppresses compartmentalization (Figure S2J). In other words, compartments are more suppressed by faster formation of the same loop architecture. We therefore hypothesized that compartmentalization is affected by the active process of loop extrusion.

Consistent with this hypothesis, recent theory and simulation studies showed that loop extrusion can alter the dynamics of a polymer as quantified by its relaxation time^102, 103^. The relaxation time, *τ*1/2(*s*), of a polymer segment is the duration of conformational “memory”, i.e. the time it takes for the segment to re-equilibrate after a perturbation (Figure 2F, S2L, and S2M; see Methods for calculation of *τ*1/2(*s*)). Extrusion events constitute such perturbations. In turn, the polymer must relax to establish compartmental contacts and spatially segregate A and B regions. In this way, extrusion may affect compartmentalization through chromosome dynamics.

Consistent with this argument, we found that compartmentalization of a segment of length *s* and its relaxation time, *τ*1/2(*s*), show the same dependence on cohesin’s residence time (Figures 2B and 2G). Initially, increasing cohesin’s residence time accelerates polymer dynamics (Figure 2G), mixing compartments and reducing compartmentalization^27^. However, further increases in residence time enable cohesin complexes to encounter and stall against each other. Such long-residing, stalled cohesin complexes are only minimally active (Figure S2N) and have little effect on chromatin dynamics; therefore, they do not disrupt contacts, and instead, they allow chromatin to relax and compartmentalize. Consistent with these arguments, compartmentalization starts recovering when *t*_res_ exceeds *τ*1/2(*s*) (Figure 2G); with short residence times chromatin cannot relax and compartmentalize, while with longer residence times, chromatin relaxes in the presence of comparatively static loops (Figure 2H). This phenomenon may also explain the recovery of near-*cis*, but not far-*cis* compartmentalization upon PDS5A/B and WAPL co-depletion (Figure 1E) because smaller genomic segments can relax more rapidly than cohesin turnover, while longer segments cannot.

Our simulations further indicate that polymer dynamics are not only influenced by cohesin’s residence time, but also by increasing extrusion velocity, which accelerates polymer dynamics (Figure 2G) and suppresses compartmentalization (Figure 2B). Here, compartmentalization is suppressed because the larger extrusion velocity increases the rate and expands the spatial extent of the disruption of contacts (Figure 2H).

Together, our polymer simulations indicate that compartmentalization is determined by the balance of conformational chromatin dynamics and loop extrusion. Interestingly, extrusion itself has a key role in determining chromatin relaxation dynamics, which further modulates compartmentalization. Thus, regulators that change cohesin’s residence time, such as WAPL and PDS5A/B, and possibly also cohesin’s extrusion speed, such as PDS5 proteins, can have profound dynamic effects on higher order chromatin organization.

### PDS5 proteins are required for the accumulation of cohesin at CTCF boundaries but not for the accumulation of CTCF at these sites

We next investigated the mechanism through which PDS5 proteins contribute to CTCF boundaries. It has been reported that depletion of PDS5B reduces CTCF binding at loop anchors^93^. We therefore tested whether simultaneous depletion of both PDS5 proteins could cause defects in CTCF boundaries by preventing the binding of CTCF to these sites. However, chromatin immunoprecipitation-sequencing (ChIP-seq) experiments did not identify clear differences in the number, genomic distribution, and signal intensities of CTCF sites between PDS5A/B depleted and control cells (Figures 3A-3C and S3A). This indicates that PDS5 proteins are not required for the association of CTCF with its binding sites and thus implies that PDS5 proteins must have other roles in the formation of CTCF boundaries.

**Figure 3.**
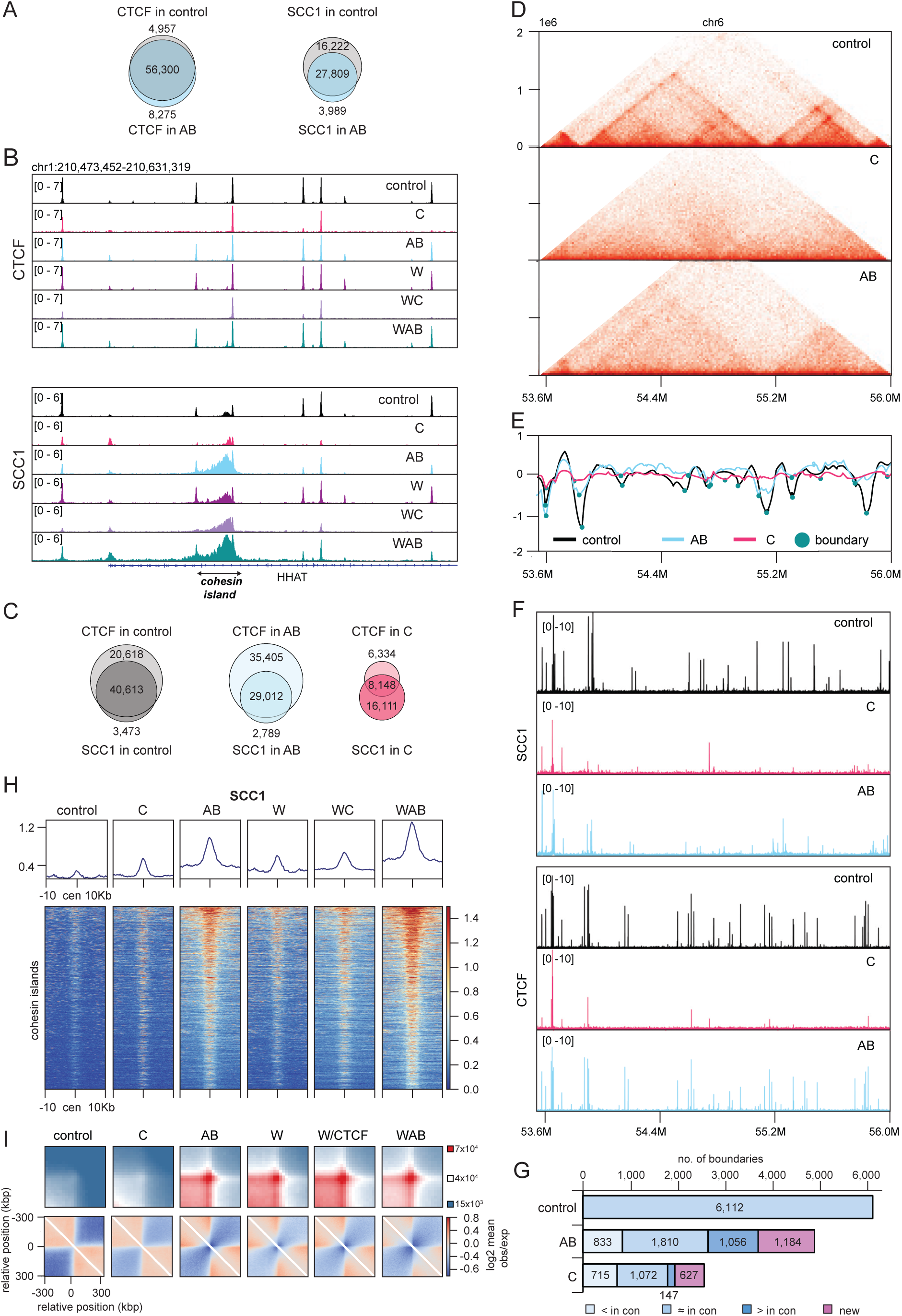
PDS5 proteins control the genomic positioning of cohesin and TAD boundaries but are dispensable for the accumulation of CTCF at its binding sites. **A.** Venn diagram illustrating the genome-wide co-localization of CTCF (left) and SCC1 (right) ChIP-seq signals from control and PDS5A/B depleted cells (AB). **B.** Binding of CTCF and SCC1 at HHAT locus, as determined by ChIP-seq in control, CTCF (C), PDS5A/B (AB), WAPL (W), WAPL/CTCF (WC) and WAPL/PDS5A/B (WAB) depletion conditions. **C.** Hi-C contact matrices for the 53.5-56 Mb region of chromosome 6 for control, CTCF (C) and PDS5A/B (AB) depleted cells. **D.** Insulation score for the same genomic region and conditions as in (C). Strong boundaries were defined by strength values bigger than 0.6 (turquoise circles). **E.** ChIP-seq profile of SCC1 and CTCF for the same genomic region and conditions as in (C). **F.** Number of strong boundaries defined as in (D) for the control, CTCF (C) and PDS5AB (AB) depleted cells and their comparison to the control boundaries. **G.** Venn diagram illustrating the genome-wide co-localization between CTCF and SCC1 ChIP-seq signals in the control (left), after PDS5A/B (AB; middle) and after CTCF (C; right) depletion. **H.** Heatmaps of SCC1 ChIP-seq signal intensities in the same conditions as in (B) around centers of detected cohesin islands +/- 10 kb, sorted by intensity over all columns; line graphs show average. **I.** *Top:* Aggregate pile-ups of Hi-C contact maps centered on pairs of neighboring cohesin islands, as identified by ChIP-seq, showing the emergence of contact peaks. Island pairs were constructed by systematically pairing each cohesin island with up to its third genomic neighbor, using a comprehensive set of detectable island regions. Coordinates are consistent across all conditions. Map resolution: 25 kb; KR-normalized. *Bottom:* Observed-over-expected pile-up at insulation boundaries in control cells.

However, in cells depleted of PDS5 proteins the accumulation of cohesin is reduced at many CTCF sites^93, 104^. Similarly, we observed in ChIP-seq experiments that the cohesin subunit SCC1 is reduced at many CTCF binding sites and instead accumulates at non-CTCF sites in PDS5A/B depleted cells (Figures 3A-3C and S3B). To test whether this reduction of cohesin at CTCF sites can explain their reduced boundary function we quantified TAD insulation genome wide^105^. This analysis identified 6,112 boundaries in control cells of which 833 (13.6%) were reduced and 3,469 (56.8%) became undetectable in PDS5A/B depleted cells, compared to 715 (11.7%) weakened and 4,325 (70.76%) undetectable boundaries in CTCF depleted cells (Figures 3D-3G). As predicted, cohesin levels were reduced at the weakened boundaries, with CTCF depletion having a stronger effect than PDS5A/B depletion (Figure 3F). These results suggest that PDS5 proteins contribute to the formation of many TAD boundaries because these proteins are required for the accumulation of cohesin at these sites, as is CTCF.

However, these analyses also revealed that the cohesin ChIP-seq and TAD insulation phenotypes of PDS5A/B depleted cells are not simply hypo-morphic versions of CTCF depleted cells. PDS5A/B depletion did not only result in the reduction of boundaries but also strengthened some boundaries (1,056; 17.3%) and caused the formation of 1,184 new boundaries that could not be detected in control cells (Figures 3G and S3C). At many of these strengthened and new boundaries, cohesin accumulated in “cohesin islands”, spanning several kb of DNA (Figures 3B, 3H, 3I, S3D and S3E), that could not be detected at the corresponding loci in control cells. About half of these cohesin islands (49.1%) were located at sites of convergent transcription (Figure S3F).

In contrast, in CTCF depleted cells only a few strengthened (147; 2.4%) and new boundaries 627) were formed and, correspondingly, hardly any cohesin islands were detected (Figures 3B, 3H, 3I and S3D). Therefore, instead of generally reducing boundaries like CTCF depletion, PDS5A/B depletion weakens many boundaries, strengthens others, and can also create new boundaries. This phenotypic difference implies that PDS5 proteins contribute to the formation or maintenance of TAD boundaries through a mechanism that is different from the one used by CTCF.

We had previously observed that cohesin islands are formed in *CTCF-Wapl* double knockout mouse embryonic fibroblasts^38^. We therefore analyzed whether cohesin islands are also formed in HeLa cells depleted of CTCF and WAPL. As expected, this was the case (Figures 3B, 3H, 3I and S3D-S3F). Interestingly, the formation of cohesin islands can therefore be induced in HeLa cells either by CTCF-WAPL depletion or by PDS5A/B depletion. This phenotypic similarity further supports the notion that PDS5 proteins control cohesin mediated loop extrusion through functions that are related to those of WAPL and CTCF.

### The role of PDS5 proteins in enabling cohesin acetylation is not sufficient to explain their role at CTCF boundaries

Since PDS5 proteins are not required for the association of CTCF with its genomic binding sites, we tested whether other functions of PDS5 proteins could explain their role in CTCF boundary formation or function. PDS5 proteins promote SMC3 acetylation^81, 84, 85^, which results in the formation of shorter cohesin loops^90, 91^. We therefore tested whether PDS5 proteins could contribute to CTCF boundary function by inhibiting loop extrusion through SMC3 acetylation.

This hypothesis predicts that the SMC3-acetylating enzymes ESCO1 and ESCO2 would also be required for TAD insulation. In previous ESCO1 depletion experiments, a partial TAD insulation defect had indeed been observed in one study^89^ but not in another^91^, but in neither of these experiments had ESCO2 been depleted. We therefore decided to compare TAD insulation and corner peaks in cells co-depleted of ESCO1 and ESCO2 (ESCO1/2) with cells co-depleted of PDS5A and PDS5B.

For technical reasons, we were unable to generate human cells lines in which both ESCO1 and ESCO2 (ESCO1/2) could be depleted. We therefore used chicken DT40 cells in which the genes encoding ESCO1 and ESCO2 had been homozygously and heterozygously deleted, respectively, and in which protein expressed from the remaining ESCO2 allele can be depleted by auxin induced degradation (*ESCO1^-/-/-^ESCO2^-/AID^*)^105^. For comparison, we also generated DT40 cells in which PDS5A and PDS5B can be simultaneously depleted by a similar strategy (*PDS5A^-/AID^ PDS5B^-/-^*; Figures S4A-S4D).

In *PDS5A^-/AID^ PDS5B^-/-^* cells treated with auxin, cohesin accumulated in vermicelli (Figure 4A), suggesting that PDS5A/B depletion increases the length of cohesin loops in DT40 cells. Hi-C experiments confirmed this interpretation by revealing that PDS5A/B depletion increased the length of *cis*-interactions genome wide (Figures 4B, 4C and S4E), whereas TAD insulation (Figure 4D), the number of corner peaks (Figure 4E), and compartmentalization (Figures S4F and S4G) were decreased. These phenotypes are similar to the ones observed in PDS5A/B depleted HeLa cells (Figures 1A-1D), indicating that the role of PDS5 proteins in CTCF boundary formation is conserved among vertebrates.

**Figure 4.**
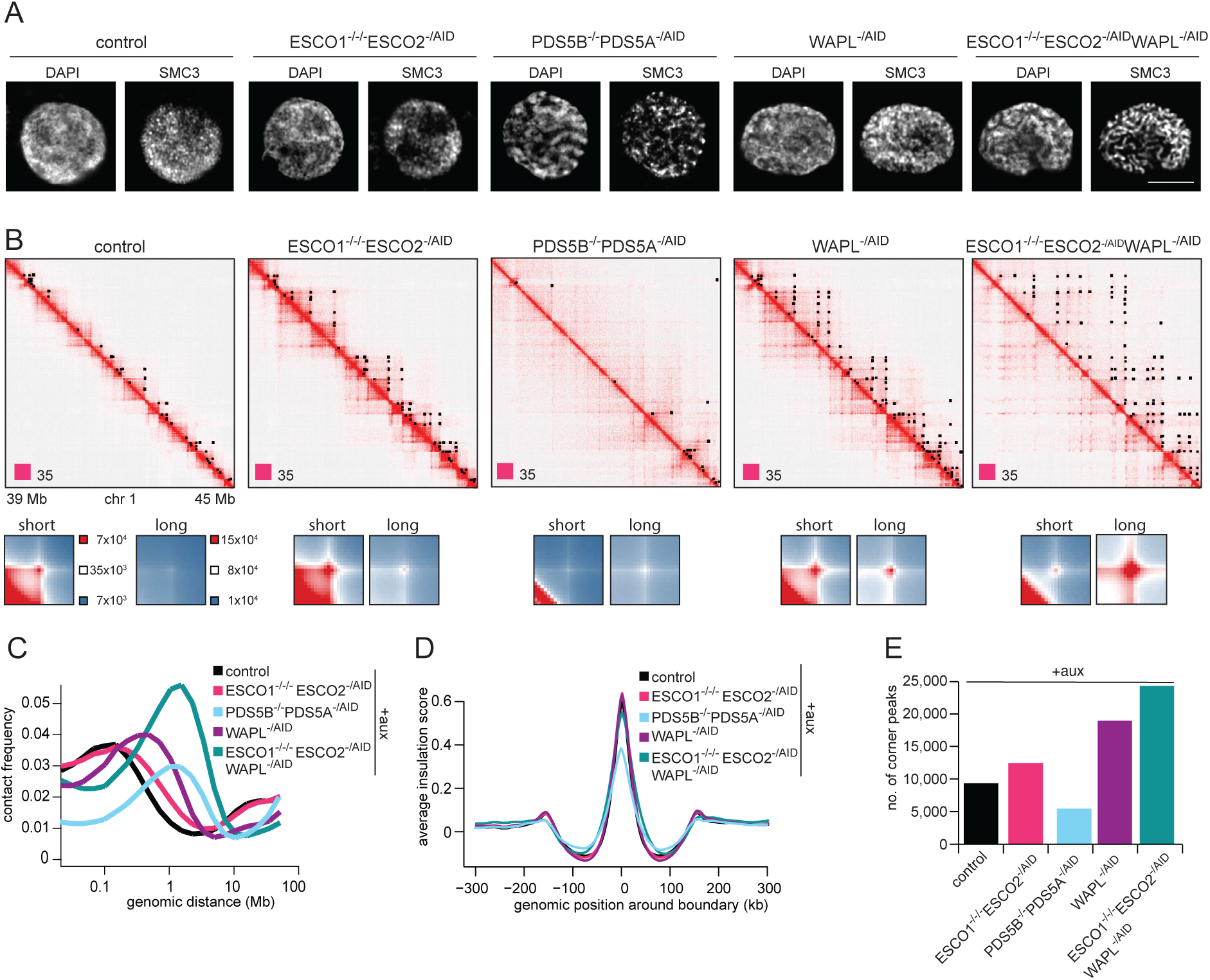
The role of PDS5 proteins in enabling cohesin acetylation is not sufficient to explain their role in CTCF boundaries. **A.** Airy scan images for Immunofluorescence staining of DAPI and SMC3-eGFP signal in the control, ESCO1/ESCO2, PDS5A/PDS5B, WAPL and triple ESCO1/ESCO2/WAPL depletion. **B.** *Top:* Coverage-corrected Hi-C contact matrices for the 39-45 Mb region of chromosome 1 plotted using Juicebox for the same condition as in (A). Corner peaks identified in this region by HICCUPS are marked by black rectangles in the upper triangle. *Bottom:* Normalized contact enrichment around corner peaks after auxin addition, for all corner peaks with 150 - 350 kb length as identified by HICCUPS in control (short) and for corner peaks above 350 kb length as identified by HICCUPS in ESCO1/ESCO2/WAPL depleted cells (long). Matrices are centered at corresponding loop anchor sites in vertical and horizontal orientation. **C.** Intra-chromosomal contact frequency distribution as a function of genomic distance for the same conditions as in (A). **D.** Average insulation score around TAD boundaries identified in control cells, shown for the same conditions as in (A). **E.** Number of corner peaks identified by HICCUPS for the same conditions as in (A).

As expected, SMC3 acetylation was reduced in PDS5A/B depleted DT40 cells (Figure S4D), consistent with the possibility that PDS5 proteins contribute to TAD insulation by enabling SMC3 acetylation. In *ESCO1^-/-/-^ ESCO2^-/AID^* cells treated with auxin, SMC3 acetylation was undetectable (Figure S4D), also as predicted. However, in these cells TAD insulation was not affected, and the number of corner peaks was not reduced but instead slightly increased (Figures 4B, 4D and 4E). These results indicate that the ability of PDS5 proteins to promote SMC3 acetylation cannot explain their role in TAD insulation and corner peak formation. It is possible that SMC3 acetylation contributes to these roles, but PDS5 must have other functions that are required for TAD insulation and corner peak formation.

These Hi-C experiments also revealed that co-depletion of ESCO1 and ESCO2 resulted in the formation of longer *cis*-interactions (Figures 4B, 4C and S4E). These results are consistent with previous ESCO1^26, 91^ and yeast Eco1 inactivation experiments^90^ and provide additional support for the hypothesis that cohesin acetylation antagonizes cohesin-mediated loop extrusion^90, 91^.

Remarkably, simultaneous inactivation of ESCO1/2 and WAPL had an additive effect on the length of *cis*-interactions and the number of corner peaks. These were both increased in *ESCO1^-/-/-^ ESCO2^-/AID^ WAPL^-/AID^*cells^105^ treated with auxin, compared to cells depleted only of ESCO1/2 (*ESCO1^-/-/-^ ESCO2^-/AID^*) or WAPL (*WAPL^-/AID^*; Figures 4B, 4C and 4SE). This additive effect can explain the previous observation that cohesin accumulates in vermicelli in these triply depleted cells but not in cells depleted of only ESCO1/2 or WAPL^105^ (Figure 4A), is consistent with results from human *ESCO1 WAPL* knockout cells^91^, and suggests that ESCO1 and ESCO2 antagonize cohesin-mediated loop extrusion by a mechanism that is different from the one used by WAPL.

### PDS5 proteins limit the amount of NIPBL bound to cohesin

Because none of PDS5A/B’s properties that we analyzed so far could explain the role of these proteins in CTCF boundaries, we tested whether PDS5 proteins perform this function by competing with NIPBL for binding to cohesin. Since NIPBL stimulates cohesin’s ATPase and loop extrusion activity but PDS5 proteins do not^63, 64, 67, 68^, such a competition mechanism could contribute to the formation of TAD boundaries and corner peaks by limiting cohesin’s loop extrusion activity, which might otherwise be able to bypass CTCF boundaries more frequently (see Discussion).

PDS5 proteins can inhibit the ATPase activity of yeast cohesin-Scc2^68^ and of human cohesin-NIPBL (Figure 5A) in a dose-dependent manner, suggesting that PDS5 proteins and Scc2/NIPBL can compete for binding to cohesin. To test this possibility directly, we performed mass photometry experiments. In these, incubation of tetrameric cohesin (542 kDa predicted molecular weight) with either PDS5B (201 kDa) or with NIPBL-MAU2 (425 kDa) led to the formation of pentameric cohesin-PDS5B (743 kDa) and hexameric cohesin-NIPBL-MAU2 complexes, respectively (997 kDa; hereafter cohesin-NIPBL; Figures 5B and S5A). When PDS5B and NIPBL-MAU2 were incubated with cohesin in equimolar amounts, both cohesin-PDS5B and cohesin-NIPBL complexes could be detected. However, when PDS5B was added in two- or fourfold molar excess, the abundance of cohesin-NIPBL complexes was reduced, whereas the abundance of cohesin-PDS5B complexes was increased (Figures 5B). PDS5B can therefore compete with NIPBL-MAU2 for binding to cohesin *in vitro*.

**Figure 5.**
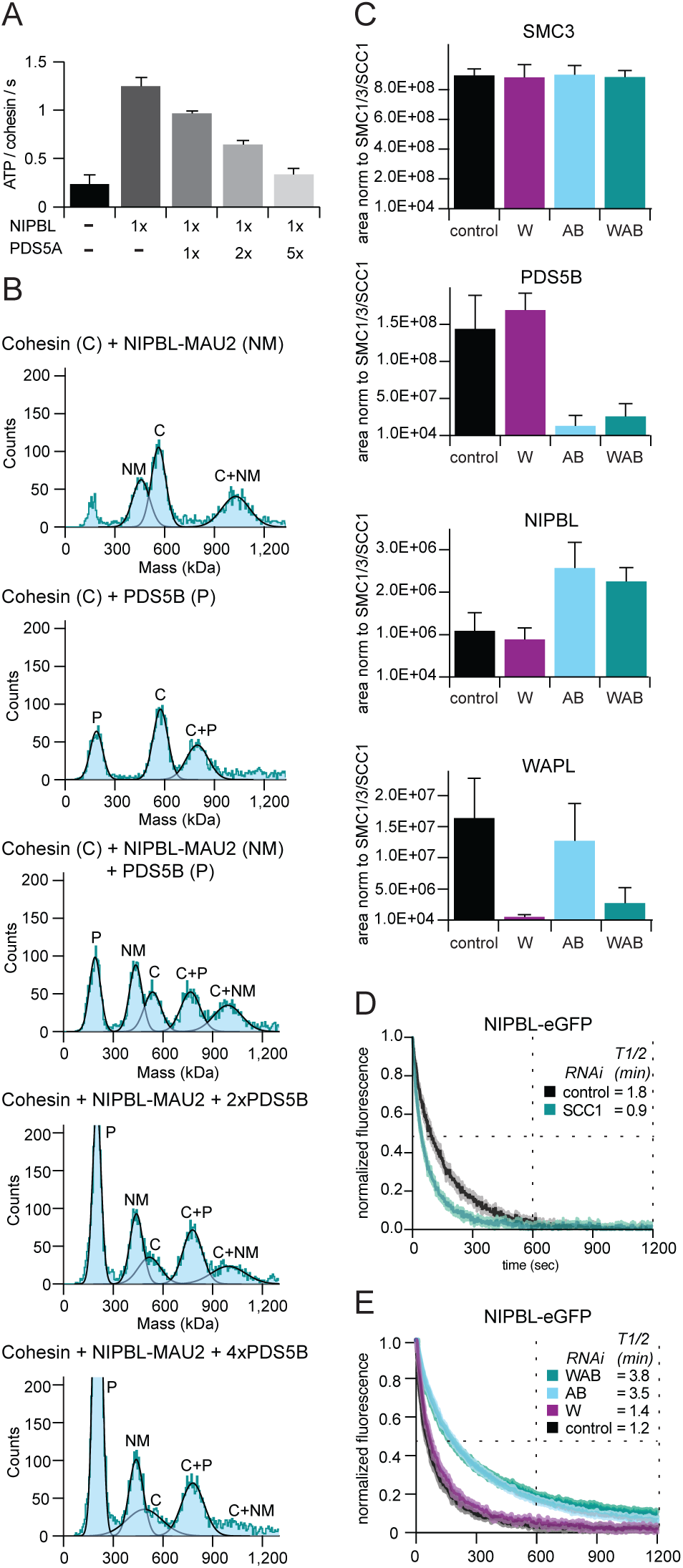
PDS5 proteins limit the amount of NIPBL bound to cohesin and the residence time of NIPBL on chromatin. **A.** ATP hydrolysis by cohesin in the presence of DNA and the indicated components (mean ± S.D. from 2 independent experiments). ‘x’ values denote the molar excess of the indicated components. **B.** Representative mass histograms (blue) and Gaussian fits of mixtures of cohesin (C), NIPBL-MAU2 (NM) and PDS5B (P). ‘x’ values denote the molar excess of PDS5B relative to cohesin and NIPBL-MAU2. **C.** Amount of NIPBL, WAPL and PDS5B bound to cohesin was determined by quantitative mass spectrometry from material immunoprecipitated using anti-GFP antibodies for SCC1- GFP cells synchronized in G1 and depleted for WAPL (W), PDS5A/B (AB) and WAPL/PDS5A/B (WAB). Peptide coverage was calculated relative to the total number of SMC3, SMC1 and SCC1 peptides. **D.** Normalised signal intensities for NIPBL- eGFP after photobleaching in control cells and cells depleted of SCC1 by RNAi. Data are presented as mean ± SD; n > 10 cells per condition. **E.** Normalised signal intensities for NIPBL- eGFP after photobleaching in control cells and cells depleted of WAPL (W), PDS5A/B (AB) and WAPL/PDS5A/B (WAB) by RNAi. Data are presented as mean ± SD; n > 10 cells per condition. Juicebox.

To test whether this is also the case in cells, we isolated chromatin-bound cohesin from PDS5A/B depleted and control HeLa cells and measured the amount of NIPBL in these samples by label-free quantitative mass spectrometry. These experiments revealed that depletion of PDS5A/B, either alone or in combination with WAPL, increased the occupancy of cohesin with NIPBL by more than two-fold (Figures 5C and S5B), suggesting that PDS5 proteins also compete with NIPBL for cohesin binding in cells. These experiments also showed that PDS5A/B depletion had little effect on the amount of WAPL that is bound to cohesin and *vice versa* (Figures 5C and S5B), further supporting the finding that WAPL and PDS5A/B can bind to cohesin separately^100^ and might thus be able to release cohesin from chromatin independently of each other (Figure 2C).

### PDS5 proteins limit the residence time of NIPBL on chromatin

To obtain further insight into how PDS5A/B regulate loop extruding cohesin through competition with NIPBL, we tested whether PDS5 proteins affect the residence time of NIPBL on chromatin. This residence time reflects, at least in part, interactions between NIPBL and chromatin-bound cohesin since cohesin depletion reduces the association of NIPBL with chromatin^69, 94^ (Figure 5D). In iFRAP experiments we observed that NIPBL has a short residence time on chromatin (t1/2 = 7 sec; Figure 5D and 5E), consistent with previous observations, which indicate that NIPBL can rapidly exchange on cohesin^63, 69, 70^. Interestingly, this residence time was increased when PDS5 proteins were depleted (t1/2 = 210 sec), either alone or in combination with WAPL, whereas WAPL depletion alone had no effect (Figure 5E).

These results are consistent with two different interpretations. PDS5A/B depletion could lengthen NIPBL’s apparent residence time by increasing the number of cohesin complexes on chromatin to which NIPBL could bind. According to this hypothesis, NIPBL would diffuse less far in PDS5A/B depleted cells before encountering a cohesin complex on chromatin that is not already bound by PDS5 proteins or another NIPBL molecule. This situation would limit the effective diffusion of NIPBL in iFRAP experiments and would be consistent with the “hopping” behavior that has been described for NIPBL^69^ (for simulations of such a scenario see^106^). Alternatively, PDS5A/B depletion could directly increase NIPBL’s residence time because PDS5 proteins have some role in dissociating NIPBL from chromatin-bound cohesin. In this case, PDS5 proteins would not simply compete with NIPBL during the cohesin binding process but would facilitate the removal of NIPBL from cohesin, i.e. increase NIPBL’s off rate.

### PDS5 proteins limit the lifetime of loop extruding cohesin-NIPBL complexes

To distinguish between these possibilities, we developed an *in vitro* three-color single-molecule imaging assay in which we could directly visualize NIPBL and PDS5 proteins during cohesin-mediated DNA loop extrusion. We labelled PDS5 proteins with JF646 and NIPBL with ATTO550 and introduced these proteins (NIPBL and either PDS5A or PDS5B) together with unlabeled cohesin-STAG1 and ATP into microscopic flow chambers. These contained lambda-phage DNA molecules that were tethered at both ends to the glass surfaces and were labelled with SYTOX Green. We then switched off the buffer flow and visualized DNA loop extrusion by total internal reflection fluorescence (TIRF) microscopy^51, 63, 70^. In most loop extrusion events (N= 239), NIPBL could be detected at the base of forming loops (91.2 %), indicating high NIPBL labeling efficiency. During the majority of these events (73 %), NIPBL became undetectable during the imaging period (∼ 9 min). In most of these cases (82.2 %), NIPBL dissociated from the DNA without detectable PDS5 binding. These NIPBL-loss events were almost always accompanied by loss of the DNA loop (98.7 %), indicating that the disappearance of NIPBL signal was caused by dissociation of NIPBL from DNA and not by photo-bleaching.

However, in a subset of cases, PDS5A or PDS5B bound to the base of the loop before NIPBL dissociated from this site (17.8 % for PDS5B; Figures 6A and S6A). In a few cases, NIPBL dissociated from the loop as soon as a PDS5 proteins could be detected on it, i.e. within one time frame (0.4 sec). Remarkably, however, in most NIPBL-PDS5 exchange events both proteins were simultaneously present on the loop for a short period of time, which, in some cases, lasted up to 11 sec (median 3 sec in case of NIPBL-PDS5B exchange events; Figure 6A and 6B), implying that NIPBL and PDS5 proteins can simultaneously bind to cohesin while it is extruding DNA.

**Figure 6.**
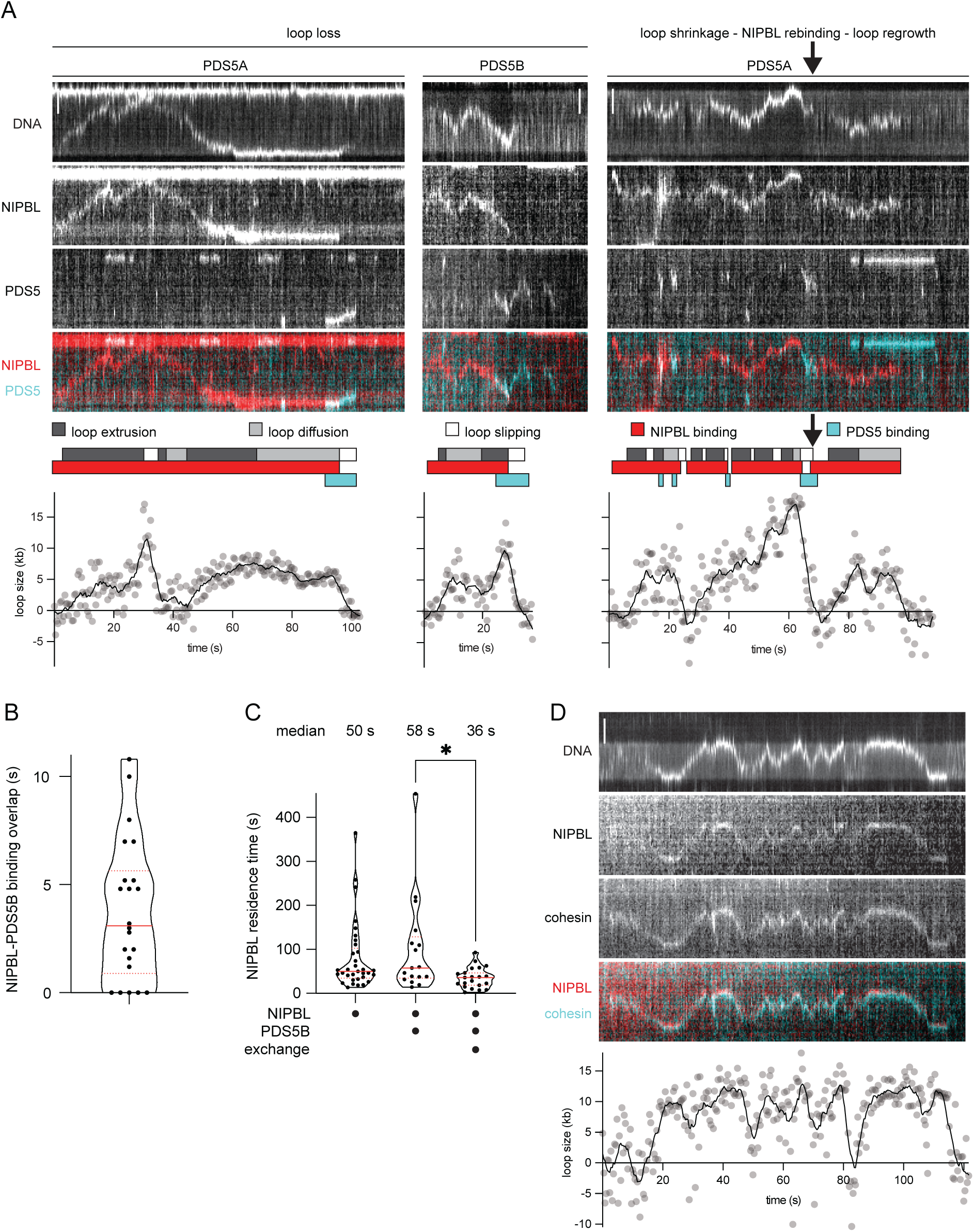
PDS5 proteins limit the lifetime of loop extruding cohesin-NIPBL complexes. **A.** Example kymographs of NIPBL-ΔN/PDS5 exchange during cohesin-mediated loop extrusion. In the left example, PDS5A binds at the loop base at 90.8 s. NIPBL-ΔN dissociates from the DNA at 95.6 s, and the DNA loop becomes undetectable at 98.8 s. In the middle example, PDS5B binds at the loop base at 24.4 s. In the right example, NIPBL-ΔN is first displaced by PDS5A and then rebinds to the loop base and displaces PDS5A.NIPBL-ΔN dissociates from the DNA at 28.8 s, and the DNA loop becomes undetectable at 32.4 s. Loop size graph dots represent raw data, solid lines represent a smoothed version using a Savitzky-Golay filter with 5 neighbors and order 0 (STAR Methods). Phases of loop extrusion, loop diffusion, loop slipping, NIPBL binding and PDS5 binding are represented by colored blocks above the loop size graph. DNA was stained with SYTOX Green. NIPBL-ΔN was labeled with ATTO550; PDS5A/B were labeled with JF646. Scale bar, 2 µm. **B.** Duration of overlap between NIPBL-ΔN and PDS5B during exchange events. Red solid line is median value. Red dotted lines are quartiles. N = 22 exchange events. **C**. Residence time of NIPBL-ΔN during loop extrusion events (left) in the absence of PDS5B, (middle) in the presence of PDS5B but without detectable PDS5B binding and (right) during loop extrusion events that end with NIPBL-ΔN-PDS5B exchange. N = 33, 17, 23 loop extrusion events. Red solid lines are median values. Red dotted lines are quartiles. Statistical significance was determined using a Mann-Whitney test (* p < 0.05). **D.** Example kymographs of NIPBL-ΔN and cohesin localization during cohesin-mediated loop extrusion. Loss of NIPBL-ΔN, cohesin and the DNA loop coincide at 116 s. DNA was stained with SYTOX Green. NIPBL-ΔN was labeled with ATTO550; cohesin was labeled with JF646.

In most NIPBL-PDS5 exchange events, the loop disappeared shortly after NIPBL had dissociated from it (Figure 6A, left and middle panels), suggesting that cohesin had unbound from at least one arm of the loop and the loop had dissolved. However, in rare cases we observed that the loop shrunk in size but did not disappear completely, implying that cohesin was still holding the loop together. In these cases, NIPBL could re-bind to the residual loop, which subsequently grew in size again (Figures 6A, right panel, arrow, and S6B).

We next tested whether PDS5 proteins have a causal role in NIPBL unbinding during these NIPBL-PDS5 exchange events, or whether these events merely represent coincidences in which the dissociation of NIPBL from a loop and the association of PDS5 occur independently from each other. To do this, we measured the residence time of NIPBL on these loops in the presence or absence of PDS5B. When we analyzed loop extrusion events in experiments in which we had not flowed PDS5B into the microscopy chambers, NIPBL had a median residence time on loops of 50 sec. This time is similar to the residence time of NIPBL on chromatin in cells^69^ (Figure 5D and E) and the time it takes *in vitro* until extruded loops fall apart once NIPBL has been washed out of the microscopy flow chambers^63^.

However, when we analyzed loop extrusion events that occurred in the presence of PDS5B, NIPBL’s residence time differed depending on whether PDS5B bound to the loop or not. In the absence of PDS5B binding events, NIPBL’s median residence time was 58 sec, i.e. similar to the time observed in the experiments without PDS5B addition. But on loops on which NIPBL was exchanged with PDS5B, NIPBL’s mean residence time was significantly shortened to 38 sec (Figure 6C). These results indicate that PDS5 proteins reduce the residence time of NIPBL on loops that are being extruded by cohesin and thus suggest that PDS5 proteins limit the lifetime of loop extruding cohesin-NIPBL complexes. Importantly, in a population of cohesin-NIPBL complexes, this effect will limit the average velocity of loop extrusion, as predicted by our simulations (Figure 2D and 2E).

As mentioned above, these results further indicate that cohesin cannot maintain a loop in the absence of NIPBL, or it can do this only for short periods of time. To test whether this is because NIPBL unbinding leads to dissociation of cohesin from DNA we imaged loop extrusion in the presence of cohesin-JF646, NIPBL-ATTO550, and DNA stained with Sytox-green. These experiments revealed that in most cases in which NIPBL dissociated from a loop when it dissolved (N = 53), cohesin also became undetectable at this site (94.4 %; Figure 6D). These results indicate that dissociation of NIPBL leads to unbinding of cohesin from the base of a loop and thus to its dissolution. This observation, combined with our finding that NIPBL-PDS5 exchange events lead in most cases to the disappearance of loops, implies that replacement of NIPBL by PDS5 proteins also results in dissociation of cohesin from DNA.

### PDS5 proteins strengthen CTCF boundaries by limiting the lifetime of cohesin-NIPBL complexes

If the ability of PDS5 proteins to limit the lifetime of cohesin-NIPBL complexes was required for the formation of CTCF boundaries, one would expect that lowering the levels of NIPBL would prevent the reduction in TAD boundaries and corner peaks observed in PDS5A/B depleted cells. To test this prediction, we generated a PDS5A/B-AID cell line in which partial NIPBL-FKBP degradation could be induced by dTAG.

When we synchronized these cells in G1 and induced degradation of NIPBL for five hours, either alone or in combination with PDS5A/B depletion, NIPBL became undetectable by Western blotting (Figure S7A). Consistent with previous NIPBL and cohesin depletion experiments^22, 24–26^, Hi-C experiments showed that *cis*-interactions, TAD insulation, and corner peak numbers were reduced in these cells, whereas compartments were strengthened (Figures S7B-S7F, S7I).

However, when NIPBL degradation was only induced for one hour, NIPBL was reduced but remained detectable (Figure S7G). Hi-C experiments revealed that this reduction in NIPBL levels indeed partially prevented the CTCF boundary defects caused by PDS5A/B depletion (Figures 7A-7H). In these experiments, we identified 12,493 corner peaks in control cells, compared to 4,054 in PDS5A/B depleted cells. However, partial co-depletion of NIPBL with PDS5A/B increased the number of corner peaks to 7,257 (Figure 7C). A partial phenotypic reversal was also observed for the length of *cis*-interactions (Figures 7A and S7H), the TAD insulation score (Figure 7B), and short-range compartmentalization (Figure S7I).

**Figure 7.**
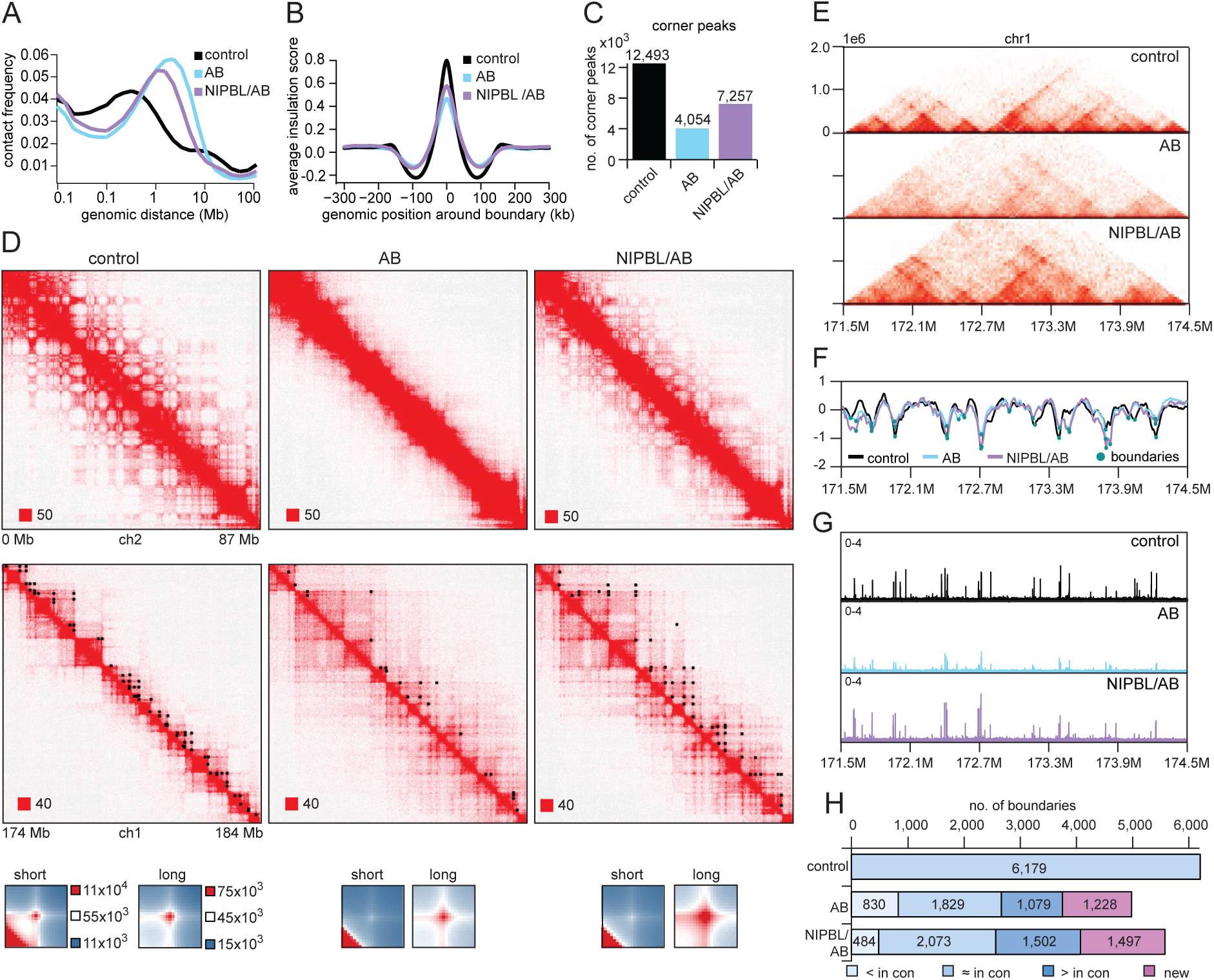
PDS5 proteins strengthen CTCF boundaries by limiting the lifetime of cohesin-NIPBL complexes. **A.** Intra-chromosomal contact frequency distribution as a function of genomic distance for control, PDS5A/B (AB), partial NIPBL over complete PDS5A/B (NIPBL/AB) depletion. **B.** Average insulation score around TAD boundaries identified in control cells for the same conditions as in (A). **C.** Number of corner peaks identified by HICCUPS for the same conditions as in (A). **D.** *Top:* Coverage-corrected Hi-C contact matrices in the 0-87 Mb of chromosome 2, plotted using Juicebox for the same condition as in (A). *Middle:* Coverage-corrected Hi-C contact matrices for the 174-184 Mb region of chromosome 1, plotted using Juicebox for the same conditions as in (A). Corner peaks identified in this region by HICCUPS are marked by black rectangles in the upper triangle. *Bottom:* Total contact counts around corner peaks in different conditions for all up to 350 kb long corner peaks identified by HICCUPS in control (short) and above 350 kb long corner peaks identified by HICCUPS in WAPL depleted cells (long). Matrices are centered at corresponding loop anchor sites in vertical and horizontal orientation. **E.** Hi-C contact matrices for the 171.5-174.5 Mb region of chromosome 1 for control, PDS5AB and cells with partial NIPBL depletion on top of complete PDS5A/B depletion (NIPBL/AB). **F.** Insulation score for the same genomic region and conditions as in (E). Strong boundaries were defined by strength values bigger than 0.6 (turquoise circles). **G.** ChIP-seq profile of SCC1 for the same genomic region and conditions as in (E). **H.** Number of strong boundaries, defined as in (F), for control cells, PDS5A/B-depleted cells (AB), and cells with partial NIPBL depletion on top of complete PDS5A/B depletion (NIPBL/AB), along with their comparison to boundaries identified in control cells.

Interestingly, the changes in TAD boundaries were more complex. In control cells, we detected 6,179 boundaries, of which 2,441 became undetectable and 830 were reduced in PDS5A/B depleted cells. Partial co-depletion of NIPBL lowered these numbers to 2,120 undetectable and 484 reduced boundaries, i.e. also in this respect caused a small reduction in the phenotype of PDS5A/B depleted cells (Figures 7F-7H). However, partial co-depletion of NIPBL with PDS5A/B did not decrease but instead increased the numbers of boundaries with increased strength (from 1,079 to 1,502) and of new boundaries (from 1,228 to 1,497; Figure 7H). Cells co-depleted of PDS5A/B and partially of NIPBL therefore contain a subset of TAD boundaries that are stronger than in control cells or not present in the control, suggesting that CTCF boundaries critically depend on the ratio of NIPBL and PDS5 proteins. ^23, 26, 94^ As expected, these changes in TAD boundaries correlated closely with changes in the genomic distribution of cohesin, which accumulated at the restored, increased, and new TAD boundaries following partial co-depletion of NIPBL with PDS5A/B (Figures 7H and S7J).

The observation that partial co-depletion of NIPBL partially restores corner peak numbers and TAD insulation in PDS5A/B depleted cells indicates that PDS5 proteins are not directly required for the formation of CTCF boundaries. Instead, these results support the hypothesis that PDS5 proteins enable the formation of CTCF boundaries by limiting the lifetime of loop-extruding cohesin-NIPBL complexes and thus the average velocity of loop extrusion.

## Discussion

DNA loop extrusion by cohesin has emerged as an important process of genome regulation^1^, and several key mechanisms have been discovered that control this activity. WAPL limits the residence time of cohesin on DNA^26, 30, 31^ and thus the processivity of cohesin-mediated loop extrusion and the length of cohesin loops^16, 22, 23, 26, 32, 33^, CTCF and other boundary elements constrain the expansion of cohesin loops in the genome^41^, and cohesin acetylation results in the formation of shorter cohesin loops, possibly by dampening cohesin’s loop extrusion activity^88, 90, 91^ (Figure 4). Here we show that PDS5 proteins control cohesin-mediated loop extrusion by a fourth mechanism, by limiting the lifetime of cohesin-NIPBL complexes. Our single-molecule visualization of NIPBL-PDS5 exchange events on loop-extruding cohesin indicates that PDS5 proteins perform this function by facilitating the dissociation of NIPBL from cohesin, and our analyses of genome architecture by Hi-C experiments and *in silico* simulations suggest that this function of PDS5 proteins has important roles at CTCF boundaries and in chromatin compartmentalization.

### PDS5 proteins limit the lifetime of loop extruding cohesin by facilitating the dissociation of NIPBL

PDS5 proteins were thought to compete with NIPBL for cohesin binding^68, 71, 91, 107^. However, our observations that binding of PDS5 to loop-extruding cohesin reduces the residence time of NIPBL on these complexes and that PDS5 depletion increases the residence time of NIPBL on chromatin in HeLa cells indicates that PDS5 proteins are not merely passive competitors, but “actively” facilitate the dissociation of NIPBL from cohesin. Consistent with this hypothesis, our results indicate that PDS5 proteins form a transient ternary complex with cohesin-NIPBL before NIPBL dissociates from cohesin. These findings imply that PDS5 proteins bind to the same site on cohesin’s kleisin subunit SCC1 as NIPBL^71^, but also, they suggest that PDS5 proteins may transiently be able to interact with other sites on cohesin. Crosslinking mass spectrometry and cryogenic electron microscopy experiments have indeed revealed additional PDS5-cohesin interactions sites at the hinge, on the ATPase head of SMC3 and on STAG1^68, 108^. It will therefore be interesting to test whether these binding sites are required for the ability of PDS5 proteins to facilitate the dissociation of NIPBL from cohesin.

It will likewise be important to understand how PDS5 proteins displace NIPBL on cohesin. Transcription factors can displace each other from single binding sites on DNA by formation of a transient ternary complex in which the incoming transcription factor facilitates dissociation of the previously bound factor^109–111^. Facilitated dissociation mechanisms have also been proposed and reported for protein-protein interaction^112–116^. Both allosteric mechanisms of facilitated dissociation and simple “invasion” mechanisms have been described, where in the latter scenario, an incoming binding partner would occlude the binding site of the previous ligand when this undergoes brief partial or complete unbinding-rebinding events^109–111, 115–117^. It is therefore conceivable that PDS5 proteins also invade the NIPBL binding site on SCC1. Alternatively, PDS5 proteins could facilitate dissociation of NIPBL allosterically by inducing conformational changes in cohesin that would trigger dissociation of NIPBL. Since cohesin undergoes large-scale conformational changes upon ATP binding and hydrolysis^118^, it will be interesting to test whether the NIPBL-PDS5 exchange can only occur during a particular step of this conformational cycle and whether PDS5 proteins promote transitions towards this step.

### What is the fate of cohesin after NIPBL-PDS5 exchange?

In our *in vitro* reconstitution experiments, most NIPBL dissociation events led to the loss of cohesin from DNA and the disappearance of extruded loops, suggesting the existence of a cohesin release reaction that does not depend on the canonical release factor WAPL. Consistent with this possibility, depletion of the NIPBL ortholog Scc2 has also been found to trigger release of cohesin from chromatin in a WAPL-independent manner during G1-phase in yeast^119^. Our finding that most NIPBL-PDS5 exchange events also lead to the disappearance of extruded loops therefore indicates that PDS5 releases loop-extruding cohesin from DNA by disrupting NIPBL-cohesin interactions.

However, since PDS5 proteins bind to acetylated cohesin with increased affinity^80, 91^ (Figure S4E) the outcome of NIPBL-PDS5 exchange reactions might be quite different when they occur on acetylated cohesin. In this case, PDS5 might remain associated with acetylated cohesin for longer periods of time, and these cohesin-PDS5 complexes might remain bound to DNA. It is further conceivable that these complexes would maintain loops that were extruded by cohesin-NIPBL and would thus “anchor” them at the genomic position at which the NIPBL-PDS5 exchange had occurred. This hypothesis will require more experimental testing, but is attractive since it could explain why PDS5 proteins and acetylated cohesin accumulate at CTCF binding sites in the genome^93, 120–122^ (Figure S3H) and how cohesin acetylation could enable the formation of long-lived loops^89, 123^.

### Can PDS5 proteins release cohesin from chromatin through two different pathways?

Previous work had shown that PDS5 proteins bind WAPL^30^, recruit WAPL to cohesin’s kleisin subunit^80^, and cooperate with WAPL in releasing cohesin from chromatin^78–82^. However, our finding that PDS5 proteins can also trigger release of loop-extruding cohesin in the absence of WAPL raises the important question whether PDS5 proteins can release cohesin from chromatin through both WAPL dependent and independent pathways. Several observations indicate that this might indeed be the case. First, a WAPL independent cohesin release pathway triggered by Scc2 depletion has been described in yeast^119^. Furthermore, our finding that co-depletion of WAPL and PDS5A/B increases cohesin’s chromatin residence time much more than depletion of WAPL alone also suggests that PDS5 proteins have WAPL-independent roles in releasing cohesin from chromatin. We therefore propose that PDS5 proteins can contribute to the release of cohesin from DNA through two distinct mechanisms, the recruitment of WAPL to cohesin, and by facilitating the dissociation of NIPBL from non-acetylated loop extruding cohesin.

### How does the ability of PDS5 proteins to facilitate the dissociation of NIPBL from cohesin contribute to CTCF boundaries?

Our finding that a partial reduction of NIPBL levels partially restores TAD boundaries and corner peak numbers in PDS5A/B depleted cells indicates that PDS5 proteins are not required for the formation of CTCF boundaries *per se*. This interpretation is further supported by our observation that Hi-C phenotypes in PDS5A/B depleted cells are not simply a hypomorphic version of the corresponding phenotypes in CTCF depleted cells. Together, these findings imply that CTCF and PDS5 proteins contribute to the accumulation of loop-extruding cohesin at CTCF boundaries in distinct ways. Our results suggest that PDS5 proteins enable efficient TAD insulation and anchoring of cohesin loops at CTCF sites, forming corner peaks, by limiting the lifetime of loop-extruding cohesin-NIPBL complexes and thereby reducing the average velocity with which loops are extruded. We suspect that this effect will strengthen the formation of TAD boundaries and corner peaks because the velocity of loop extrusion will determine how frequently cohesin-NIPBL complexes arrive at CTCF sites, and this frequency will determine how often loop-extruding complexes bypass these CTCF sites. *In silico* simulations have indicated that CTCF boundaries are dynamic because CTCF can frequently unbind and rebind^124^. This model implies that the frequency with which CTCF boundaries are bypassed increases with the frequency with which loop-extruding complexes arrive at these sites because more encounters with these sites afford more opportunities for bypassing an unoccupied site. CTCF boundaries therefore do not only control cohesin loops, but cohesin-mediated loop extrusion conversely also regulates the efficiency with which these boundaries function.

These results and the fact that PDS5 proteins are essential for mouse embryogenesis^73^ imply that the levels of PDS5 proteins might be particularly relevant in situations in which the ability of CTCF sites to stop loop-extruding cohesin is functionally important, such as during V(D)J recombination^32, 60^ and protocadherin promoter choice^62^. It will therefore be interesting to analyze whether PDS5 proteins are regulated to modulate the velocity of loop extrusion and the efficacy of CTCF boundaries in particular physiological situations, analogous to how WAPL levels are regulated in developing B cells and neurons^16, 32, 33^. In this context it is curious to note that the *A. thaliana* genome encodes five PDS5 paralogs ^125^, suggesting that cohesin-NIPBL’s lifetime might be subject to more elaborate control mechanisms in plants.

### How do PDS5 proteins control chromatin compartmentalization?

It has been proposed that loop extrusion interferes with chromatin compartmentalization by actively mixing chromatin^27, 29, 126^ and physically bridging different compartments^25, 27^. In addition, the formation of vermicelli in PDS5A/B or WAPL-depleted cells^31^ was hypothesized to prevent chromosome folding into compartments due to the stiffness of the bottlebrush polymer structures^27^. In contrast, we find that loop extrusion primarily impacts chromatin compartmentalization through dynamics, while effects of the loop architecture itself are more subtle.

In simulations, cohesin extruding through a chromosomal segment disrupts existing chromatin contacts, such as those formed by compartmentalization, because extrusion continuously forms new contacts. If relaxation of the chromosomal segment is slow (*τ*1/2(*s*)>*t*_res_), such a disruption locally interferes with equilibration, i.e., the ability to establish homotypic interactions and spatially segregate A and B regions. However, if the segment relaxes rapidly (*τ*1/2(*s*)<*t*_res_), cohesin complexes with long residence times on chromatin may facilitate compartmentalization. This competition between extrusion and polymer equilibration determines (Figure 2H): 1) the ability of the chromosomal segment to compartmentalize and 2) whether perturbations to extrusion are suppressive or stabilizing for compartmentalization. Our findings are consistent with recent theoretical work showing that 3D chromatin compaction depends on the polymer relaxation time compared to the residence time^102, 103^. These principles could be experimentally tested by measuring chromatin relaxation times in two-point locus-tracking experiments^56^ with fluorescent tags separated by different genomic distances. Measurement of these timescales would enable a direct comparison to cohesin’s residence times, which, in parallel with Hi-C experiments could test the predicted effects on compartmentalization.

Together, our results indicate that PDS5 proteins have indirect, but profound effects on compartmentalization by decreasing the residence time and velocity of loop-extruding cohesin complexes. PDS5 proteins intermittently displace NIPBL from cohesin, thus stalling and dissociating cohesin, which limits the duration, rate, and extent of the disruption of compartmental contacts. Although depletion of PDS5 proteins leads to the accumulation of cohesin in vermicelli, vermicelli formation does not inherently prevent compartmentalization (Figure 2A-B). Instead, PDS5A/B depletion shifts extrusion into a regime of longer residence times and faster extrusion, leading to longer durations of larger disruptions of compartmental contacts, due to comparatively slow polymer relaxation (Figure 2D, 2E, and 2G). In contrast, the extremely long cohesin residence times resulting from PDS5A/B and WAPL co-depletion generate nearly static chromosome bottlebrushes with stalled cohesin complexes and very long-lived loops, allowing relaxation and compartmentalization. Thus, PDS5 proteins directly alter cohesin dynamics, but have variable downstream effects on chromatin compartmentalization, depending on the presence of other regulators of extrusion dynamics and chromatin conformational dynamics. Altogether, we find that molecular regulation of cohesin governs both extrusion-mediated contacts via cohesin-NIPBL activities and mesoscale compartmentalization via polymer dynamics.

## Limitations of study

Our conclusion that NIPBL-PDS5 exchange events lead to dissociation of cohesin from DNA is based on the observation that these exchange events result in the loss of DNA loops, but we could not visualize NIPBL, PDS5, cohesin and DNA simultaneously since our microscopy system can only detect three colors. We have also not yet reconstituted NIPBL-PDS5 exchange events on acetylated cohesin and do therefore not know whether PDS5 remains bound to acetylated cohesin for longer periods of time and whether these cohesin-PDS5 complexes remain bound to DNA. Finally, our experiments do not reveal where in the genome NIPBL-PDS5 exchange events take place. We suspect that these events can take place anywhere in the genome, but since both acetylated cohesin and PDS5 proteins accumulate at CTCF sites ^89, 93, 120–122, 127^ (Figure S3H) it is conceivable that NIPBL-PDS5 exchange events stabilize acetylated cohesin-PDS5 complexes at these sites.

## Acknowledgements

We thank Dr. Takuya Abe for help with imaging DT40 cells, VBCF for next generation sequencing, IMP proteomics facility for mass spectrometry and Elphege Nora and Geoff Fudenberg for communicating unpublished results. We thank Viraat Goel and Anders Hansen, as well as Timothy Földes, Max Imakaev, Henrik Pinholt, and the other members of the Mirny Lab for helpful discussions. Research in the laboratory of J-MP is supported by Boehringer Ingelheim, the European Union ((ERC); AdG 101020558 LoopMechRegFun, MSCA 101072505 CohesiNet) and the Vienna Science and Technology Fund (10.4739/LS19-029). DB is supported by a research grant from the Italian Association for Cancer Research (AIRC) – Project-ID 30385. LAM and EJB are supported by NSF (MCB2044895 and ANR 2210558), LAM is a Simons Investigator (Simons Foundation International). For the purpose of Open Access, the authors have applied a CC BY public copyright license to any Author Accepted Manuscript (AAM) version arising from this submission. J-MP is also an adjunct professor at the Medical University of Vienna and an external member of the Max Planck Institute of Biochemistry, Martinsried.

## Author contributions

GW, IFD, and J-MP designed the experiments and interpreted the data. RS and LC bioinformatically analyzed and interpreted data. WT generated HeLa cell lines. RK generated DT40 cell lines. WT and GW performed ChIP-seq experiments. GW generated Hi-C libraries, performed Western blotting and immunoprecipitation experiments. KN and GW carried out and analyzed iFRAP experiments. IFD performed single-molecule experiments. EJB and LM designed the simulation studies. EJB performed and analyzed *in silico* simulations. RJ acquired Airyscan images and developed the script for vermicelli quantification. The project was supervised by LM, DB, and J-MP. The manuscript was written by GW, IFD, EJB, LM, and JMP.

## Material and methods

### Cell line generation

The HeLa Kyoto (female) cell lines used in this were generated in the following way. The CRISPR/Cas9 mediated genome editing cell lines SCC1-GFP and Halo-AID-WAPL/SCC1-GFP were created as described previously^26, 94^. Similarly mKate2-AID-PDS5A/Halo-AID-PDS5B/SCC1-EGFP, Halo-AID-WAPL/mKate2-AID-PDS5A/SNAP-AID-PDS5B/SCC1-EGFP, CTCF-AID-mKate2/Halo-AID-WAPL/SCC1-EGFP and HA-FKBP12^F36V^-NIPBL/mKate2-AID-PDS5A/Halo-AID-PDS5B/SCC1-EGFP were generated using the following gRNAs (PDS5A: CACCGTGCGCGGTGAAGTCCATCC and CACCGTGGCGTCGTGAGTGCCG AC; PDS5B: CACCGCTGATATTTCCTTGACCCC and CACCGCAAAGACTAGGACCAATGA; CTCF: CACCGCAGCATGATGGACCGGTGA and CACCGGAGGATCATCTCGGGCGT G). Clones were selected after verification of homozygous integration by PCR of genomic DNA (primers used for PDS5A: CAAGGGCGTTTTGTTCCCG and ACCTTCTGTCAAAGTTTAACCACC; PDS5B: GTTGGTGGGGAAGGTTACCA and ACCTCGATCGGAAATGCTCAA; CTCF: ACTGTTAATGTGGGCGGGTT and CTGGGTGCCATTCTGCTACA). Based on the cell line SCC1-Halo-P2A-Tir1^128^, we introduced AID-mKate2 tag to the C-terminus of CTCF, generating CTCF-AID-mKate2/SCC1-Halo-P2A-Tir1. gRNA sequences and genotyping primers were described above.

For auxin mediated degradation in DT40 cells we engineered a DT40 cell line with a deletion in the catalytic domain of ESCO1. The ESCO1 gene is on chromosome 2 which is present in three copies, thus resulting in an ESCO1-/-/- cell line. Subsequently, we generated ESCO2-/3AID6FLAG conditional cells in this background, allowing for the down-regulation of ESCO2 upon auxin addition ^105^. PDS5A-/3AID6FLAG PDS5B-/- cells were generated in the following way (Fig S4A). PDS5B-KO vectors were generated from genomic PCR products combined with Puromycin and Ecogpt selection marker cassettes. Left arm and right arm of PDS5B-KO vectors were amplified using the primers GATCTCATGACAGTCTGTGAGTTAC and GAACTGGGGGCTTTTTGTATC; and GTGAGGGAGATACTGTGTCACAGG and GTCAGATGTGCAGTGTGGGATTCC (for the right arm of the KO construct). Amplified PCR products were cloned into pLoxP vectors^129^. The PDS5B-KO vectors were linearized by KpnI before the transfection. To construct PDS5A-KO vector, genomic PCR products were combined with Neomycin marker cassette. Left arm and right arm were amplified using the primers TCCACACTGCATTAATCCGTAGGTG and CAATTCTCTGCGATGAACAGGGAC; and GCAGAGAGTTGCATAGGGTTGCATG and TACACAGGGCTGTAGAGACCTTTGG (for the right arm of the KO construct). Amplified PCR products were cloned into pLoxP vectors. The PDS5A-KO vector was linearized by PvuI before the transfection. PDS5A-AID vector was created by combining the homology arm amplified by primers GTCTTTCACATACAGGCAGCACTTC and TTGTAGATCGATCTGTCTCTGTGCTGC, and p3xmAID-6xFLAG vector with L-histidinol marker cassette ^130^. The PDS5A-AID vector was linearized by SacI before the transfection. To validate the insertion of the AID-tag at the C-terminus of PDS5A, genomic PCR products were generated with a pair of primers that are designed upstream of the homology arm and in the middle of the targeting construct. For the validation of PDS5B gene disruption, two primers designed on the deletion locus and downstream of the 3’-homology arm. A primer set amplifying PARP1 gene locus was used as a control.

### Cell synchronization

To analyse the consequences of deffernet depletions in HeLa cells, cells were synchronised at early S phase by two consecutive rounds of treatment with 2 mM <u>thymidine</u> (Sigma). Eleven hours after second thymidine release, cells were treated with 200 μM auxin (Andole-3-Acetic Acid, Gold Biotechnology) for 5h and then collected. DT40 ells were synchronized by nocodazole treatment for 6 hours, followed by release into a medium containing mimosine and auxin for 8 hours. At this stage, most cells were in the late G1/early S phase, as confirmed by fluorescence-activated cell-sorting (FACS) analysis (Fig S1A).

### Immunofluorescence

Cells were fixed with 4% formaldehyde for 20 min at room temperature and permeabilized with 0.1% Triton X-100 in PBS for 5 min. After blocking with 3% BSA in PBS-T 0.01% for 30 min, the cells were incubated with the primary antibodies for 1 h and subsequently incubated with the secondary antibodies together with DAPI for 1 h at room temperature. Finally, the coverslips were mounted with ProLong Gold (Thermo Fisher) before imaging.

### RNA interference

For RNAi experiments, the cells were transfected as described previously^131^. Briefly, the cells were transfected by incubating 100 nM duplex siRNA with RNAi-MAX transfection reagent in antibiotic-free growth medium. After 48 h of RNAi treatment, cells were harvested for experiments. The following target sequences of siRNAs (Ambion) were used: SCC1 (5′- GGUGAAAAUGGCAUUACGGtt -3′); WAPL (5’-CGGACUACCCUUAGCACAAtt-3’); PDS5A (5′- GCUCCAUAUACUUCCCAUGtt -3′); PDS5B (5’- GAGACGACUCUGAUCUUGUtt -3’)

### iFRAP

The cells used for iFRAP were previously engineered to express a live-cell CDK2 sensor^132^, allowing the distinction of cells in the G1 phase from those in other cell cycle stages. Cells were seeded on chambered coverglass (Nunc 155409) for 2 days in DMEM supplemented with 10% FCS, 0.2 mM L-glutamine antibiotics. After washing with pre-warmed cell culture medium three times, cells were incubated for 30 min and then exchanged for pre-warmed phenol red free medium supplemented with 10% FCS, 0.2 mM L-glutamine antibiotics for imaging. Live cell imaging was performed using an LSM880 confocal microscope (Carl Zeiss), equipped with a 40x /1.4 numerical aperture (N/A) oil DIC Plan-Apochromat objective at 37°Cand 5 % CO2. Photobleaching half of the nucleus was performed by 2 iterations of a 561 nm diode laser at max intensity after the acquisition of two images. After the bleaching, images were acquired at 1 min intervals for 2 hours.

### Chromatin fractionation and immunoprecipitation

For chromatin fractionation, cell pellets were extracted in a buffer consisting of 20 mM Tris (pH 7.5), 150 mM NaCl, 5 mM MgCl2, 2 mM NaF, 10% glycerol, 0.2% NP40, 20 mM b-glycerophosphate, 0.5 mM DTT, and protease inhibitor cocktail. Chromatin pellets were separated from whole cell extracts by centrifugation at 2,000 g for 5 min and were washed three times with the same buffer. Genomic DNAs in the resulting chromatin pellets and in whole cell extracts were digested in the same buffer supplemented with Benzonase nuclease (homemade). For chromatin immunoprecipitation, cells were cultured in five 600 cm2 trays (Thermo, cat#166508) were arrested for 20-24 hours with 2 mM Thymidine and released into early G phase for 10 hours. Auxin was added for 5h after which the cells were, scraped off, washed with cold PBS, and snap-frozen. The pellets were lysed by syringing through 22G needles in 2-3 volumes of lysis buffer (75 mM HEPES pH 7.5, 1.5 mM MgCl2, 150 mM KCl, 15% glycerol, 1.5 mM EGTA pH 8.0, 0.075% NP40, 1 mM DTT, 0.1 mM PMSF, 10 μg/ml leupeptin, 10 μg/ml chymostatin, 10 μg/ml pepstatin, 20 mM b-glycerophosphate, 10 mM NaF, 1 mM Na3VO4). Chromatin bound fraction was separated from soluble by centrifugation at 1,500 g for 5 min at 4°C. The chromatin pellet was washed 5 times by re-suspension in 10 ml Wash buffer I (50 mM HEPES pH 7.5, 1.5 mM MgCl2, 150 mM KCl, 10% glycerol, 1.5 mM EGTA pH 8.0, 0.075% NP40, 1 mM DTT, 0.1 mM PMSF, 10 μg/ml leupeptin, 10 μg/ml chymostatin, 10 μg/ml pepstatin, 20 mM b-glycerophosphate, 10 mM NaF, 1 mM Na3VO4). The final chromatin pellet was re-suspended in 2 volumes of Wash buffer I, supplied with 300 U/ml Benzonase, and rotated for 2 hours at 4°C. The reaction was then centrifuged for 2 min at 2,500 g and the supernatant, containing released chromatin-bound proteins, was used for IP. IP was performed by a 2-hour rotation at 4°C with 100 μl ChromoTek GFP-Trap® Magnetic Agarose Beads. Beads were then washed 4 times with 10 volumes Wash buffer I, 7 times with Wash buffer II (50 mM Hepes pH 7.5, 1.5 mM MgCl2, 150 mM KCl, 10% glycerol, 1.5 mM EGTA pH 8.0), and once with 150 mM NaCl. Beads with the bound protein were submitted for Mass spectrometry.

### Calibrated ChIP-Seq

An equal number of cells from each condition were mixed with 3 % MEFs expressing EGFP-SMC3 and then fixed with 1 % formaldehyde for 10 min at room temperature. After quenching by 125 mM Tris-HCl pH 7.5, cells were washed with PBS and snap frozen in liquid nitrogen. Subsequent ChIP-seq experiment procedures were performed as described ^133^. Briefly, cell pellets were thawed on ice and lysed with lysis buffer (50 mM Tris-HCi pH 8.0, 10 mM EDTA pH 8.0, 1 % SDS, 1 mM PMSF and protease inhibitor cocktail). Cell lysates were sonicated for 5 cycles (30 sec on/off) at maximum power using a Biorupter to shear the genomic DNA. 10x volumes of dilution buffer (50 mM Tris-HCi pH 8.0, 10 mM EDTA pH 8.0, 1 % Triton X-100, 1 mM PMSF and proteasome inhibitor cocktail) were added to lysate. After pre-clearing by incubation with Affi-Prep protein A beads, the lysates were incubated with antibodies overnight at 4°C and incubated with the same beads for 3 hours at 4°C. The beads were washed 2 times with wash buffer 1 (20 mM Tris-HCi pH 8.0, 2 mM EDTA pH 8.0, 1 % Triton X-100, 150 mM NaCl, 0.1 % SDS and 1 mM PMSF), wash buffer 2 (20 mM Tris-HCi pH 8.0, 2 mM EDTA pH 8.0, 1 % Triton X-100, 500 mM NaCl, 0.1 % SDS and 1 mM PMSF), wash buffer 3 (10 mM Tris-HCi pH 8.0, 2 mM EDTA pH 8.0, 250 mM LiCl, 0.5 % NP-40, 0.5 % deoxycholate) and TE buffer (10 mM Tris-HCl pH 8.0, 1 mM EDTA pH 8.0), and then eluted with elution buffer (25 mM Tris-HCi pH 7.5, 5 mM EDTA pH 8.0 and 0.5 % SDS) for 20 min at 65°C two times. The elutes were treated with RNase-A at 37°C for 1 hour and then proteinase K at 65°C overnight. DNA was purified by phenol/chloroform/isoamyl alcohol (25:24:1) extraction and ethanol precipitation. DNA was resuspended in 100 μl H2O, and ChIP efficiency was quantified by quontitative PCR (qPCR). Sequencing libraries were prepared using the NEBNext® Ultra II kit following the manufacturer’s instructions, and DNA samples were submitted for Illumina deep sequencing at the Vienna Biocenter Core Facilities.

### Hi-C library preparation

We generated a total of 51 in situ Hi-C libraries from our experiments (Table S1). Libraries were prepared using the MboI restriction enzyme, all HeLa samples, were generated following the protocol described by Rao et al. (2014) without modification. Briefly, the in situ Hi-C protocol involves crosslinking cells with formaldehyde, permeabilizing nuclei with detergent, digesting DNA overnight with a 4-cutter restriction enzyme, filling in 5′ overhangs while incorporating a biotinylated nucleotide, ligating the resulting blunt ends, shearing the DNA, capturing biotinylated ligation junctions with streptavidin beads, and analyzing the resulting fragments by paired-end sequencing. For DT40 cells, all libraries were generated using the 6-base cutter HindIII. The same Hi-C protocol was applied, except that cells were lysed by homogenization with pestle A (2 × 30 strokes) after a 15-minute incubation in ice-cold lysis buffer. Chromatin was solubilized in 0.5% SDS at 65 °C for 7 minutes. In all cases, NEB adapters were ligated using the NEBNext Ultra II DNA Library Preparation Kit for Illumina.

### ATPase assay

Recombinant cohesin was incubated as indicated with NIPBL-MAU2 and PDS5A in a 25 µl reaction (final assay conditions: 25 mM NaH2PO4/Na2HPO4 pH 7.5, 35 mM NaCl, 2.5 mM MgCl2, 1 mM DTT, 0.1 mg/ml BSA, 2 mM ATP, 10 nM [γ-^32^P]ATP (Hartmann Analytic; SCP-501), 10 ng/µl λ-DNA, 60 nM cohesin, 60 nM NIPBL-MAU2, 60/120/300 nM PDS5A). Reactions were incubated at 37 °C and stopped by adding 1 % SDS and 10 mM EDTA. Reaction products were separated on polyethyleneimide plates (Sigma; 1055790001) by thin-layer-chromatography using 0.75 M KH2PO4 (pH 3.4), analyzed by phosphor imaging with a Typhoon Scanner (GE Healthcare) and quantified with ImageJ.

### Mass photometry

Recombinant cohesin, NIPBL-MAU2 and PDS5B were incubated on ice as indicated in a 10 µl reaction for 5 min (reaction conditions accounting for carry-over from protein storage buffers: 7.6 mM NaH2PO4/Na2HPO4 [pH 7.5], 45 mM NaCl, 1.5 % glycerol, 15 mM imidazole [pH 7.5], 32 mM Tris [pH 7.5], 32 mM potassium glutamate, 2 mM MgCl2, 2 mM ATP, 100 nM cohesin, 100 nM NIPBL-MAU2, 100/200/400 nM PDS5B). 2 µl of this reaction was added to the gasket containing 18 µl of imaging buffer (50 mM Tris [pH 7.5], 50 mM potassium glutamate, 2.5 mM MgCl2) for subsequent mass photometry measurement (see below).

Mass photometry data was recorded using a OneMP (Refeyn Ltd, Oxford). Coverslips (24 × 50 mm, Marienfeld, VWR 630-2187) were cleaned by sequential sonication in Milli-Q water for 5 min and then in isopropanol for 10 min. Coverslips were then rinsed extensively with Milli-Q water, sonicated in Milli-Q water for 5 min and dried using a nitrogen stream. Measurements were performed in silicon gaskets (GBL103250, Grace Bio-Labs) attached to precleaned coverslips. Prior to the addition of the protein mixture (see above), 18 µl of imaging buffer (50 mM Tris [pH 7.5], 50 mM potassium glutamate, 2.5 mM MgCl2) was added to the gasket and the focus position was adjusted to the glass-liquid interface. 2 µl of protein mixture was then added to the gasket and a 60 second movie was recorded using AcquireMP software (Refeyn Ltd.). Movie processing was performed using DiscoverMP software (Refeyn, Ltd.) and measured contrasts were converted to mass as described^66^.

### Protein expression and purification

#### Recombinant cohesin and NIPBL-MAU2

All steps were performed at 4 °C unless indicated. For cohesin (c357), NIPBL-MAU2 (LC50A) and NIPBL-ΔN (BB241) purified from Sf9 insect cells, around 20 ml cell pellets were lysed using Dounce homogenization in purification buffer 1 (25 mM NaH2PO4/Na2HPO4 [pH 7.5], 500 mM NaCl, 5% glycerol) supplemented with 10 mM imidazole [pH 7.5], 0.05 % Tween 20, 1 mM PMSF, 3 mM beta-mercaptoethanol, 10 µg/ml aprotinin, 2 mM benzamidine (Sigma), and cOmplete EDTA-free protease inhibitor cocktail. Following centrifugation at 48,000 g for 45 minutes, the supernatant was mixed with 5 ml of Toyopearl AF-chelate-650M resin (Tosoh Bioscience) pre-charged with Ni^2+^ ions and incubated for 3 hours. The beads were washed with 3 x 10 bead volumes of purification buffer 1 supplemented with 15 mM imidazole [pH 7.5] and 0.01% Tween 20. Protein was eluted with 25 ml of purification buffer 2 (25 mM NaH2PO4/Na2HPO4 [pH 7.5], 150 mM NaCl, 5% glycerol, 300 mM imidazole [pH 7.5], 0.01% Tween 20), combined with 5 ml of FLAG-M2 agarose (Sigma; A2220) and incubated for three hours. The beads were washed with 3 x 10 bead volumes of purification buffer 3 (25 mM NaH2PO4/Na2HPO4 [pH 7.5], 150 mM NaCl, 5% glycerol, 50 mM imidazole [pH 7.5]). Protein was eluted by 3 x 10 min incubations with 1 bead volume of purification buffer 3 supplemented with 0.5 mg/ml 3xFLAG peptide (MDYKDHDGDYKDHDIDYKDDDDK). Eluates were concentrated to around 0.5 ml using Vivaspin 20 (50 kDa MWCO) ultrafiltration units (Sigma; GE28-9323-62), frozen in liquid nitrogen, and stored at -80 C.

#### HeLa cohesin

All steps were performed at 4 °C unless indicated. Around 5 ml cell pellet was thawed and resuspended in 40 ml of buffer A (20 mM Tris [pH 7.5], 1.5 mM MgCl2, 10 mM KCl) supplemented with 0.5 mM DTT, 1 mM PMSF, 10 µg/ml aprotinin and 1x cOmplete EDTA-free protease inhibitor cocktail (Roche; 11873580001). Cells were lysed using 10 strokes of Dounce homogenization, incubated on ice for 10 minutes, and then homogenized for another 15 strokes. Nuclei were pelleted by centrifugation at 2000 rpm for 15 minutes. Pelleted nuclei were resuspended in 36 ml of buffer A supplemented with 1 mM PMSF and 1x cOmplete EDTA-free protease inhibitor cocktail and homogenized with 3 strokes. NaCl was added to a final concentration of 500 mM dropwise while stirring. Tween 20 was then added to 0.1%, and the lysate was stirred for 10 minutes before sonication (Branson Digital Sonifier; 60 x 0.5 s pulses at 40% amplitude). Following centrifugation at 48,000 g for 30 minutes, the soluble fraction was combined with 1 ml of FLAG-M2 agarose resin and incubated for three hours. Beads were washed with 3 x 10 bead volumes of buffer B (25 mM NaH2PO4/Na2HPO4 [pH 7.5], 150 mM NaCl, 5% glycerol, 1 mM EDTA). Bound protein was eluted with 5 ml of buffer B supplemented with 0.5 mg/ml 3xFLAG peptide. The eluate was concentrated to approximately 0.2 ml using a Vivaspin 2 100 kDa MWCO ultrafiltration unit (Sigma; GE28-9322-57), frozen in liquid nitrogen, and stored at -80 °C.

#### PDS5A and PDS5B

All steps were performed at 4 °C unless indicated otherwise. Sf9 cells (around 10 ml cell pellet) were thawed and lysed by Dounce homogenization in purification buffer 1 (25 mM NaH2PO4/Na2HPO4 [pH 7.5], 500 mM NaCl, 5% glycerol) supplemented with 10 mM imidazole [pH 7.5], 0.05 % Tween 20, 1 mM PMSF, 3 mM beta-mercaptoethanol, 10 µg/ml aprotinin, 2 mM benzamidine (Sigma), and cOmplete EDTA-free protease inhibitor cocktail. After centrifugation (48,000 g, 45 min), the soluble fraction was combined with 1 ml of NiNTA agarose (Qiagen; 30230) and incubated for 90 min. Beads were washed with 2 x 50 ml purification buffer 1 supplemented with 20 mM imidazole [pH 7.5] and 0.01 % Tween 20 and 1 x 50 ml PDS5 wash buffer (25 mM NaH2PO4/Na2HPO4 [pH 7.5], 150 mM NaCl, 5% glycerol) supplemented with 20 mM imidazole [pH 7.5]. Bound protein was eluted with 12 ml PDS5 wash buffer supplemented with 300 mM imidazole [pH 7.5]. The eluate was concentrated to around 0.6 ml using a Vivaspin 6 (50 kDa MWCO) ultrafiltration unit (Sigma; GE28-9323-18), filtered through a 0.22 µm spin column (Millipore; UFC30GV00) and applied to a Superdex 200 10/300 GL column at 0.4 ml/min equilibrated in size exclusion buffer (25 mM NaH2PO4 / Na2HPO4 [pH 7.5], 150 mM NaCl, 5% glycerol, 1 mM DTT). Fractions containing PDS5 were pooled, concentrated using a Vivaspin 6 (50 kDa MWCO) ultrafiltration unit (Sigma; GE28-9323-18), frozen in liquid nitrogen and stored at -80°C.

### Fluorescent labeling of HeLa cohesin, NIPBL-ΔN, PDS5A and PDS5B

ATTO550-HaloTag and JF646-HaloTag ligands were generated as described^70^. To label HeLa cohesin, following a 3 h incubation of the nuclear extract with FLAG-M2 agarose resin (see above), the beads were washed with 2 x 10 bead volumes of buffer B (25 mM NaH2PO4/Na2HPO4 [pH 7.5], 150 mM NaCl, 5% glycerol, 1 mM EDTA). Beads were resuspended to one bead volume in buffer B, supplemented with excess JF646-HaloTag ligand and incubated for 20 min at room temperature protected from light. Beads were washed extensively with buffer B and protein was eluted, concentrated and frozen as described above. NIPBL-ΔN was labeled with ATTO550-HaloTag ligand as for HeLa cohesin, except beads were washed with purification buffer 3 (25 mM NaH2PO4/Na2HPO4 [pH 7.5], 150 mM NaCl, 5% glycerol, 50 mM imidazole [pH 7.5]). PDS5A and PDS5B were labeled using the same approach, except labeling was performed while proteins were bound to Qiagen NiNTA resin, and beads were then washed with PDS5 wash buffer (25 mM NaH2PO4/Na2HPO4 [pH 7.5], 150 mM NaCl, 5% glycerol) supplemented with 20 mM imidazole, followed by elution and size exclusion chromatography as described above.

### Loop extrusion assay

DNA loop extrusion experiments were performed essentially as described^1, 51^. Glass surfaces were pegylated with a mixture of 5 mg/ml methoxy-PEG-N-hydroxysuccinimide (MW 3500, LaysanBio) and 0.05 mg/ml biotin-PEG-N-hydroxysuccinimide (MW 3400, LaysanBio) in 0.1 M NaHCO3, 0.55 M K2SO4 buffer^70, 134^. Surfaces were pegylated a further four times with MS(PEG)4 (MW 333.33, Thermo Fisher) in 0.1 M NaHCO3. Slides and coverslips were incubated at 4 °C overnight in the dark, washed with Milli-Q water and dried using a nitrogen stream.

Flow chambers were assembled by sandwiching a pegylated coverslip with a drilled glass slide using an adhesive sheet (Grace Bio-Labs; SA-S-1L) and sealed using epoxy glue. Inlet and outlets were connected to PE60 tubing (Avantor Sciences; 63019-070). Avidin DN (Vector Laboratories) was diluted to 0.2 mg/ml in buffer 1 (20 mM Tris [pH 7.5], 150 mM NaCl, 0.25 mg/ml BSA (Thermo Fisher Scientific; AM2618) and drawn into the flow chamber using a 1 ml syringe and incubated for 15 min. The flow chamber was washed with 1 ml of buffer 1 and then transferred to the incubation chamber of a Zeiss Elyra 7 with Lattice SIM^2^ microscope pre-warmed to 37 °C. From this point onwards, all buffers and reaction mixtures were pre-warmed to 37 °C prior to introduction into the flow chamber. The outlet was connected to a syringe pump (Harvard Apparatus; 70-4504) and was washed with 0.4 ml buffer 1 supplemented with an additional 1 mg/ml BSA. End-biotinylated lambda-DNA (48.5 kb)^135^,was diluted to 6 pM in buffer 1 supplemented with 10 nM SYTOX Green (Thermo Fisher Scientific; S7020) and introduced into the flow chamber at 6 µl/min such that DNA molecules were stretched to a length of about 5 µm (about 30% of their contour length). The flow chamber was washed first with 0.2 ml of buffer 1 supplemented with 10 nM SYTOX Green, then with 0.4 ml of buffer 2 (50 mM Tris [pH 7.5], 200 mM NaCl, 1 mM MgCl2, 5 % glycerol, 0.25 mg/ml BSA), then with 0.4 ml of imaging buffer (50 mM Tris [pH 7.5], 50 mM potassium glutamate, 2.5 mM MgCl2, 0.25 mg/ml BSA) supplemented with 10 nM SYTOX Green and finally with 0.1 ml imaging buffer supplemented with oxygen scavenger mix (final concentrations: 0.2 mg/ml glucose oxidase (Sigma; G2133), 35 µg/ml catalase (Sigma; C-40), 9 mg/ml D-glucose, 2 mM trolox (Cayman Chemical, 10011659)), 10 nM SYTOX Green and 5 mM ATP ( Jena Biosciences, NU-1010-SOL). For NIPBL/PDS5A/B dual labeling experiments, HeLa cohesin was introduced into the flow chamber at 1.1 nM, NIPBL-ΔN-ATTO550 at 0.75 nM and PDS5A/B-JF646 at 0.75 – 2.25 nM in imaging buffer supplemented as above. For cohesin/NIPBL dual labeling experiments, HeLa cohesin-JF646 was introduced into the flow chamber at 0.6 nM and NIPBL-ΔN-ATTO550 at 0.6 nM. 60 µl of protein mixture was flowed into the flow chamber at 100 µl/min. Time-lapse microscopy images were then acquired using a Zeiss Elyra 7 with Lattice SIM^2^ equipped with 488 nm, 561 nm and 639 nm lasers, two PCO Edge 4.2 sCMOS cameras and a ×63/1.46 NA Alpha Plan-Apochromat oil objective. All data were acquired in HiLO mode using an exposure time of 100 ms per channel; images were acquired sequentially such that the time interval for each channel was 400 ms. Imaging was performed for 9 min following flow in of proteins.

### Loop extrusion assay image analysis

All loop extrusion events in a single field of view in the absence of buffer flow were manually identified, and kymographs were generated for each channel in Fiji using 9-pixel wide straight line selections and the Multi Kymograph command. For the examples presented in Figures 6 and S7, kymographs were adjusted for brightness and contrast only. To determine the size of DNA loops, we used two custom scripts written in the ImageJ macro language. First, a bandpass filter was applied to reduce noise and enhance the signal. Semi-automated tracking was performed via finding local maxima within the search radius and manual alignment was used when none was found. Mean intensities at the position of loop extrusion and in the background were measured for each frame. In a second step, a straight-line selection was drawn to capture a kymograph for further analysis. The background was subtracted per frame and sum intensities were measured for a 9-pixel diameter circular selection at the position of loop extrusion, as well as for the remaining non-looped DNA on both sides of the circular selection. The amount of DNA within the loop at each timepoint (*Loop*) was calculated by dividing the intensity at the position of loop extrusion by the total intensity of all DNA segments combined. Since the 9-pixel diameter circular selection included non-looped DNA, the *Loop* values were subtracted from 9-pixel diameter circular selection intensity values averaged over 12 frames prior to the onset of loop extrusion. These corrected *Loop* values were converted to kb by multiplying by 0.48502, i.e. the total length of bacteriophage lambda genomic DNA, and plotted as a function of time using Graphpad Prism.

#### NIPBLΔN characterisation and NIPBL/PDS5 exchange quantification

Kymographs of all loop extrusion events in a single field of view were generated as described above. The residence time of NIPBLΔN in these loop extrusion kymographs was defined as the period between the first timepoint at which NIPBLΔN was detected until its disappearance. For NIPBL/PDS5 exchange quantification, loop extrusion events where either 1. no NIPBL signal was detectable or 2. PDS5A/B was detectable on DNA throughout the imaging period were excluded from further analysis. For the NIPBLΔN residence time analysis presented in Figure 6 and S6, NIPBLΔN dissociation events were manually classified into two groups: 1. NIPBLΔN dissociation events that occurred in the absence of PDS5A/B binding; 2. NIPBLΔN dissociation events that occurred in the presence of PDS5A/B binding. Residence times of NIPBLΔN were calculated as described above.

## QUANTIFICATION AND STATISTICAL ANALYSIS

### Imaging of interphase nuclei with ZEISS LSM 880 (Airyscan) and image quantification

To achieve high resolution of cohesin staining upon different depletions, advanced confocal microscopy by using the Airyscan system in combination with a ZEISS LSM880 was applied. Images were acquired with a 63x plan-apochromat lens (*NA* = 1.40) in immersion oil (refractive index *n* = 1.514). For each channel, appropriate lasers for excitation (λex) and matching filters (λem) were chosen. DAPI: λex = 405 nm, 0.2% laser, 850 digital gain, λem = 440 nm; Alexa-488 (SCC1): λex = 488 nm, 0.1% laser, 850 digital gain, λem = 515 nm. Using Airyscan SR mode, z stacks with a 100 nm spacing and 15 slices were recorded with an image dimension of 45 µm, 35 nm pixel size and a pixel dwell time of 0.82 µs. Stacks were centred according to the approximate centre of each nucleus in z.

The acquired raw images were reconstructed and deconvolved with Huygens Professional 21.04 (SVI) by using SuperXY mode with subsequent correction of xy shift and rotation relative to the z axis.

For quantification of the vermicelli phenotype, a Fiji macro script was written. The script measures the vermicelli area within a nucleus as well as the total area of the nucleus to calculate the proportion of the nuclear area that is occupied by SCC11 signal, or in other words the degree of compaction. Vermicelli were selected and the area was measured in the SCC1 channel according to auto-thresholding with the Otsu algorithm and exclusion of any signal outside the nucleus. The nuclear area of each cell was measured according to Otsu thresholding of the DAPI signal.

### iFRAP curve fitting

The fluorescent recovery kinetics of SCC1-eGFP and NIPBL-eGFP after photobleaching were quantified using the ZEN2011 software, as the difference between the mean fluorescence signal intensity of the bleached and the unbleached regions followed by background subtraction. pciFRAP curves were normalised to the mean of the pre-bleach fluorescent intensity and to the first image after photobleaching. G1 cells were identified by nuclear or cytoplasmic localization of DHB-mVenus signals, respectively ^132^. Individual iFRAP curves were fitted using the single exponential function f(t)=EXP(-kOff1^∗^t) or the double exponential function f(t)=a^∗^EXP(-kOff1^∗^t)+(1-a)^∗^EXP(-kOff2^∗^t) in R using the mini-pack.lm package (version 1.2.1) as previously described^136^. Double exponential curve fitting was performed under the constraints that 1/kOFF1 (the dynamic residence time) and 1/kOFF2 (the stable residence time) were in the range of 1-40 min and 5-15 hours, respectively. The statistical details can be found in the figure legends.

### ChIP-Seq peak calling and calibration

Prior to treatments and Illumina short read sequencing, a fixed percentage of mouse cells expressing SMC3 LAP containing GFP marker protein was added to all samples as a calibration reference throughout the entire follow-up processing. Sequencing reads which passed the Illumina quality filtering were mapped against a fusion genome <u>template</u> constructed of mm9 (MGSCv37) and hg19 (GRCh37) reference assemblies. The subsets of uniquely mappable reads from the mm9 resp. hg19 fractions (allowing up to two mismatches each) were separated for further steps. The mm9 fractions from all samples were processed using <u>peak calling</u> by MACS version 1.4.3 with the same fixed shift-size parameters resulting in a genome-wide set of peak regions for the Flag-tagged reference protein in all reference mouse cell populations^137^. Considering the expected evenness in signal for all samples in a hypothetical loss-free procedure, out of all peak-sets a union peak set was built with constant coordinates being common in all sample references. This common coordinate set’s read abundances throughout all samples delivers reliable measures for usage in overall calibration of the various treated human hg19 fractions. Peak callings of the actual human (hg19) samples with MACS software versions 1.4 and 2 were applied on full, merged, filtered, and deduplicated replicate data sets using background input (-dox). As a significance threshold we applied a p-value of 1e-10 resulting in reliable peak sets which are confirmed also through very low numbers of detectable negative peaks in the range of few dozen from -dox cells. Calibration factors described above were applied to coverage values of alignment profile tracks of the samples to overcome the problem of experimental variation in yield efficiency. The statistical details can be found in the figure legends.

### Calculation of peak overlaps

Genomic overlaps have been calculated using multovl version 1.3^138^ and area-proportional threefold Venn diagrams were drawn with eulerAPE^139^. Since occasionally more than one site from one dataset overlaps with a single site in a second dataset, the resulting coordinates of such an overlap contribute to one single entry - a so-called union. Consequently, the overall sum of site counts drops slightly if displayed in union overlaps. The statistical details can be found in the figure legends.

### Identification of cohesin islands

Cohesin islands were identified as areas of clustered peak appearance, as described before^38^. In brief, if individual narrow peaks fulfilling the p-value threshold of 1e-10 are positioned sufficiently close to each other to overlap within the range of half of their own size, then they get merged. If resulting multiply merged peak regions exceed 5 kb in size, then they were categorized as a cohesin island. The statistical details can be found in the figure legends.

### Hi-C data processing

Illumina sequencing was performed on all Hi-C libraries with 150 bp paired-end reads. Two replicate data sets for each library were truncated, filtered, and aligned against the human genome assembly hg19 (GRCh37) using bowtie2 and the HiCUP processing pipeline version 0.7.4, and finally merged ^140^.

Alignments were converted into input for juicer_tools pre (juicer tools version 1.22.016; as well as input for HOMER version 4.11 (http://homer.ucsd.edu/homer/.98 PMID: 20513432). All juicer-based contact matrices used for analysis were Knight-Ruiz normalized.

Loop annotation, also known as corner peak annotation, and merging was performed using ‘juicer_tools hiccups’ at default resolution. TAD annotation and insulation scores were generated by the ‘findTADsAndLoops.pl’ script in the HOMER software package. This software scans the relative contact matrices for locally dense regions of contacts or areas with an increased degree of intra-domain interactions relative to surrounding regions. Plots of insulation profiles were made using insulation scores in the bedGraph file format and TAD boundary coordinates from regions around these coordinates. Aggregate peak analysis of hiccups-called Hi-C peaks was performed using ‘juicer_tools apa’. The Hi-C looping peaks detected from all the conditions were classified according to their sizes and used as coordinates for aggregation within Hi-C maps. These aggregated size-classified Hi-C peaks were visualized by plotting the cumulative stack of sub-matrices in a way that Hi-C peaks lie at the center of the matrix and that the resulting APA plot displays the abundance of contacts respective to local contact density.

### Pearson correlation matrix

FAN-C (https://fan-c.readthedocs.io/en/latest/index.html) was used to produce the Pearson correlation matrix at 500 kb resolution, shown in 1E (bottom). The compartment switches in Figure 1G were calculated using an eigenvector analysis performed with Juicer-tools (https://github.com/aidenlab/juicer) at a 250 kb resolution.

Pile-up analysis in Figure 3I (bottom) was done with cooltools (https://github.com/open2c/cooltools) using the merged coordinates of cohesin islands identified in the indicated genotypes.

Figure 3c-d-e were generated using the open2c tools (https://github.com/open2c/open2c_examples) using a 100 kb window sliding at a 5 kb resolution. The threshold for boundaries was defined at 0.6. The graph in 3F uses the previously calculated strong boundaries coordinates in each condition. We compared the boundary strength of each condition to the control and used plus/minus 0.2 threshold to define them. Finally, the boundaries with a strength higher than control were further divided into new boundaries or overlapping with control using a 20 kb window on each side.

### Compartment contrast and contact frequency

Hi-C reads with mapping quality ≥30 were mapped to 1 kb resolution cooler files^141^ using the hg19 genome assembly for Hela cells and the galGal6 assembly for DT40 cells via the distiller v0.3.4 pipeline in the default setup (https://github.com/open2c/distiller-nf/tree/v0.3.4) ^142^ to create cooler files. For Hela, computations below were performed using chromosomes 1-4, 7, 9, 10, 12-15, 17, 18, 20-22, and X; computations for DT40 used chromosomes 1-28 and 30-33. Maps were balanced by genome-wide iterative correction^143^.

Contact frequency as a function of genomic separation [*P*(*s*)] was computed as the sum of *cis* contacts at each genomic distance (diagonal), *s*, scaled by the number of valid pixels at that distance. *P*(*s*) was averaged over chromosomes and smoothed. The calculation was performed using the expected_cis method from the cooltools library^144^. Log derivatives, 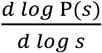, were computed using the numpy gradient function.

Compartments were assigned by computing the first eigenvector at 50 kb resolution using the eigs_cis method from cooltools^144^, correlating with GC content to determine eigenvector orientation^145^; eigenvectors were visually inspected for accuracy.

Compartment contrast ^27^ at each genomic distance, *s*, was computed as the difference between the sum of intra-compartmental (*i.e.*, AA and BB) contact frequencies and inter-compartmental contact frequencies (*i.e.*, AB), scaled by the sum of intra- and inter-compartmental contact frequencies:

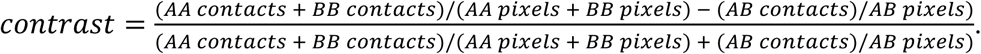

The result was smoothed with gaussian_filter1d function from scipy.ndimage, with sigma=2.

### Polymer simulations

We performed polymer molecular dynamics simulations using freely and publicly available code from previous work^146^ with new custom configuration files for three repeats of a 20 Mb segment (207-227 Mb) of Hela chromosome 1 (see https://github.com/mirnylab/microcompartments/tree/main/comp_extr). This code uses the polychrom v0.1.0 library^147^, which is a Python wrapper for the OpenMM molecular simulation toolkit^148, 149^.

Simulations couple 1D loop extrusion activity on the chromatin polymer to 3D polymer structure and dynamics. Pairs of genomic sites occupied by a loop extruder at an instant in time are physically bridged, therefore holding a polymer loop. The polymer has A and B compartments, which microphase separate via affinity interactions. Compartment assignments are determined from experimental data from Hela cells. Between loop extrusion simulation steps, the polymer with loops evolves via Langevin dynamics.

#### Loop extrusion simulations

We simulate *N* loop extruders (i.e., cohesins) on a 1D lattice of *L*=60000 genomic sites, where each site represents σ=1 kb of chromatin. Unless noted, *N=*120 such that we have mean separation *d*=500 kb. At each time step, each loop extruder may unload chromatin, after which it immediately rebinds, and each component of the two components of each loop extruder may perform a translocation step. A loop extruder is loaded at a pair of adjacent unoccupied sites. Loop extrusion occurs as each of the two components of the extruder translocate away from the position at which it was loaded. Unless specifically varied or otherwise noted, translocation occurs at speed *v*=1 kb/s^63, 64, 150^, which achieved by setting the stepping probability per extrusion time step to *p*=0.5. Translocation by an extruder component continues until: a) the loop extruder is unloaded, which occurs stochastically with characteristic residence time *t*_res_, which was set to 100 s except in the specific sweeps varying *t*_res_ or b) it encounters an obstacle, such as another loop extruder or the end of the polymer. For simulations with frozen loops (as in Figures S2K), we first performed 1D active extrusion simulations performed for 10^6^ extrusion steps to equilibrate the loop distribution. Loop extruders were then frozen in place on the polymer chain. These frozen extruders maintained fixed polymer loops throughout the subsequent 3D polymer simulation.

#### Polymer dynamics

We simulate *L*=60000 monomeric subunits, each of diameter *a≈*25 nm and representing σ=1 kb of chromatin, connected by harmonic springs to form a linear polymer. Subunits also interact by a soft repulsive potential, which models excluded volume interactions. We assigned A- and B-type monomers, based on compartment assignments from experiments. B-type monomeric subunits interacted attractively with affinity *εB*=0.05 *kBT* via a smooth square well potential, as in previous studies^27, 146^. The polymer was confined at 20% volume fraction^151^ within a sphere. Polymer dynamics were computed using the OpenMM fixed timestep Langevin integrator with (polychrom variables) timestep=40 and collision_rate=0.01. Polymer simulations were evolved for 1100 timesteps between each loop extrusion timestep, except for simulations with *v* > 2 kb/s, which used 550 polymer timesteps per extrusion timestep. Loop extruder configurations from the 1D extrusion simulations were used to set the anchors of polymer loops, which were bridged together, at the beginning of each block of polymer timesteps. Choices for monomer diameter and polymer steps per extrusion step resulted in simulation time and length comparable to previous simulations^96, 146^ and experiments in mammalian cells^56^.

#### Contact maps and analysis

Data was collected for a minimum of 10 independently equilibrated simulations per condition. Simulation contact maps at 4 kb resolution were generated with the contactmaps.binnedContactMap method from polychrom^142^ using a contact radius of 4 monomer diameters. Contact maps were computed by time and ensemble averaging the 3 chromosomal repeats and the independent simulations. Compartment contrast was computed as in the experimental analysis.

#### Computation of chromatin polymer relaxation time

Conformational memory is quantified by the time-correlation function, *Cs*(*t*), of the end-to-end vectors, 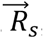, for polymer segments of different sizes, *s*, within the chromosome. The correlation function is defined as 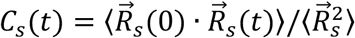, where 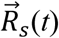 is the vector between the two ends of a polymer segment of length *s* at time *t* and 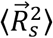 is the mean-squared end-to-end distance of a segment of length *s*. The correlation function decays over time, with a characteristic half-life, *τ*1/2(*s*), referred to as the relaxation time^102, 103, 152^. Equivalently, the relaxation time defined in this manner is equal to the average time required for the two-point mean-squared displacement, 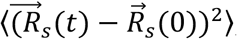, to surpass the mean-squared end-to-end distance, 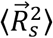 (*i.e.*, the time for the two ends of the polymer segment to explore the region occupied by the segment; Figures S2L and S2M).

## Supplemental figures

**Figure S1.**
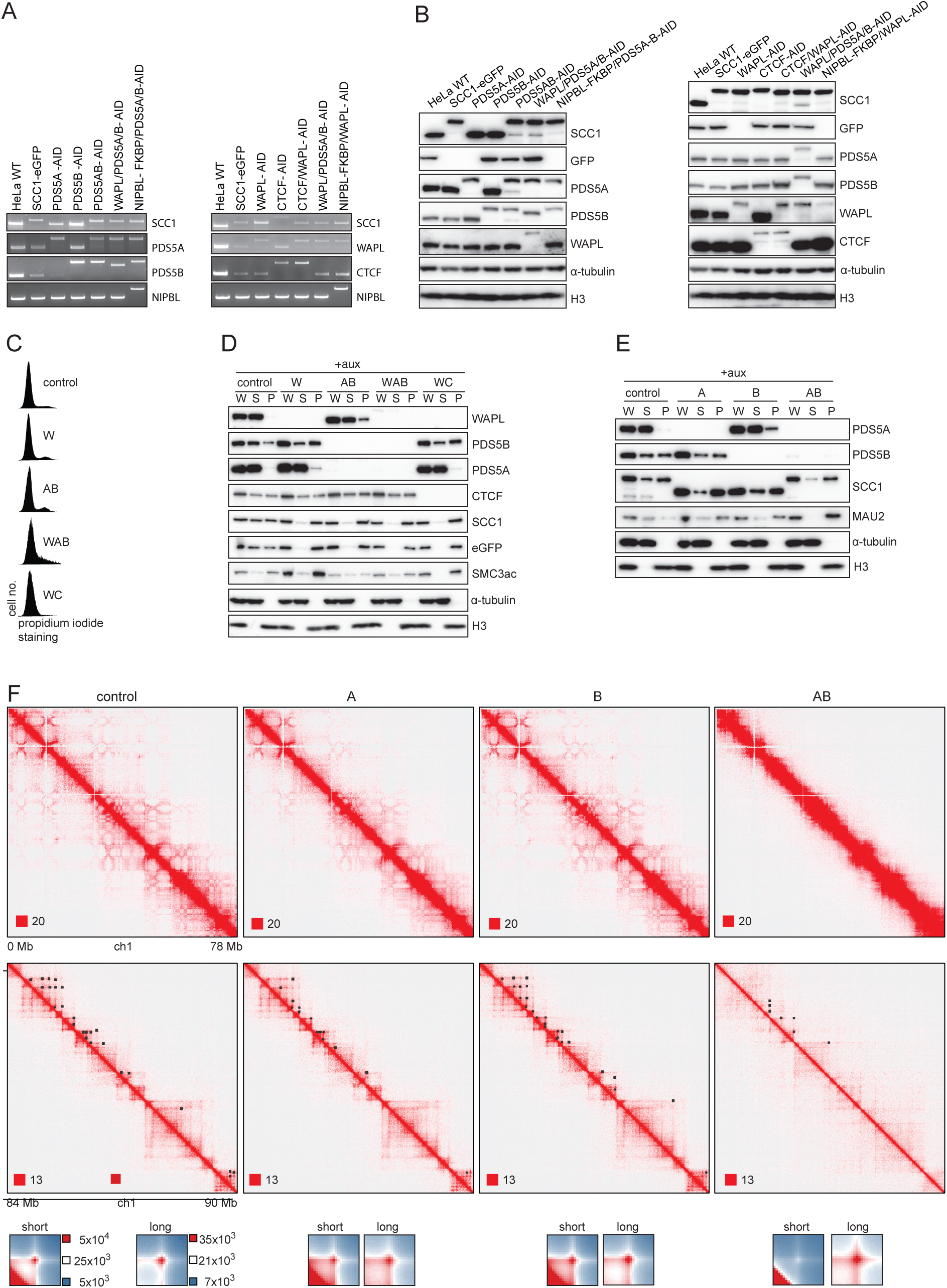
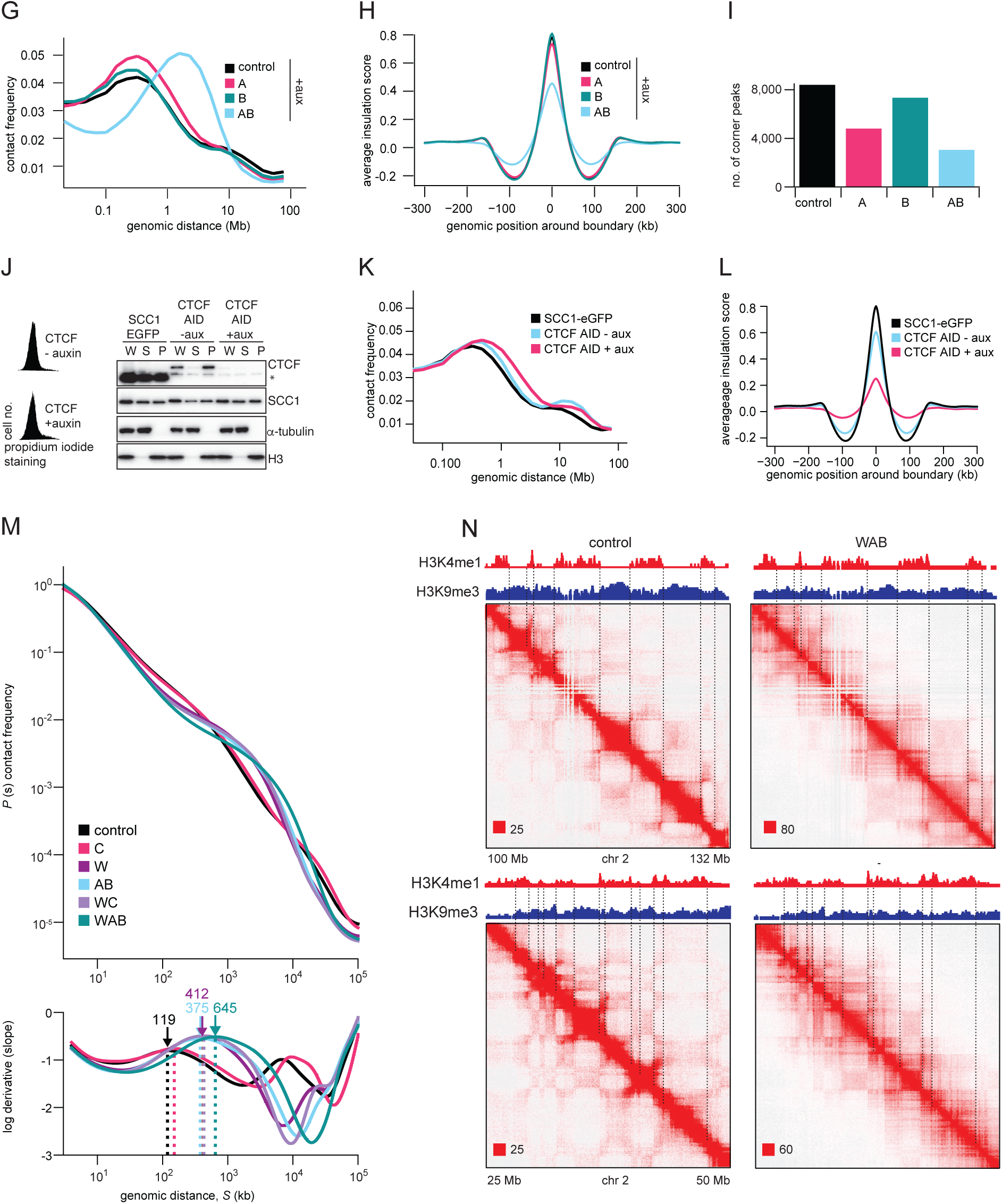
Characterization of degron cell lines and additional Hi-C phenotypes, related to Figure 1. **A.** Genomic PCR genotype analysis of the indicated cell lines. Left panel, from left to right: parental HeLa WT, SCC1-eGFP, mKate2-AID-PDS5A (PDS5A-AID), Halo-AID-PDS5B (PDS5B-AID), mKate2-AID-PDS5A/Halo-AID-PDS5B/SCC1-eGFP (PDS5A/B-AID), Halo-AID-WAPL/mKate2-AID-PDS5A/SNAP-AID-PDS5B/SCC1-eGFP (WAPL/PDS5A/B-AID) and HA-FK-BP-NIPBL/mKate2-AID-PDS5A/Halo-AID-PDS5B/SCC1-eGFP (NIPBL-FKBP/PDS5AB-AID). Right panel, from left to right: parental HeLa WT, SCC1-eGFP, Halo-AID-WAPL/SCC1-eGFP (WAPL-AID), CTCF-AID-mKate2/SCC1-Halo-P2A-Tir1 (CTCF-AID), CTCF-AID-mKate2/Halo-AID-WAPL/SCC1-EGFP (CTCF/WAPL-AID), Halo-AID-WAPL/mKate2-AID-PDS5A/SNAP-AID-PDS5B/SCC1-eGFP (WAPL/PDS5A/B-AID) and HA-FKBP-NIPBL/Halo-AID-WAPL/SCC1-eGFP (NIPBL-FKBP/WAPL-AID). Genomic PCR products were generated with primers that annealed outside of the homology arm that was used for inserting different tags. **B.** Immunoblotting analysis of whole-cell extracts isolated from cell lines described in (A). α-Tubulin and H3 were used as loading controls. **C.** Flow-cytometry profiles of control cells, cells depleted for WAPL (W), PDS5AB (AB), WAPL/PDS5A/B (WAB) and WAPL/CTCF (WC) synchronized in G1 by double thymidine block. **D.** Chromatin fractionation and immunoblot analysis of auxin-treated SCC1-mEGFP (control), PDS5AB-AID (AB), WAPL/PDS5A/B-AID (WAB) and WAPL/CTCF-AID (WC) cells expressing Tir1. After auxin addition, whole-cell extract (W), supernatant (S) and chromatin pellet fractions (P) were analyzed by immunoblotting using antibodies against the proteins indicated on the right. α-Tubulin and H3 were used as loading controls. **E.** As (D) except for SCC1-eGFP (control), PDS5A-AID (A), PDS5B-AID (B) and PDS5A/B-AID (AB) cells expressing Tir1. **F.** Top and middle: coverage-corrected Hi-C contact matrices for the 0-78 Mb (upper panels) and 84-90 Mb (middle panels) regions of chromosome 1 plotted using Juicebox for control, PDS5A, PDS5B and double PDS5AB depleted cells. Bottom: Normalized contact enrichment around corner peaks after auxin addition, for all corner peaks with 150 - 350 kb length as identified by HICCUPS in control cells (short) and above 350 kb long corner peaks identified by HICCUPS in WAPL depleted cells (long). Matrices are centered at corresponding loop anchor sites in vertical and horizontal orientation. **G.** Intra-chromosomal contact frequency distribution as a function of genomic distance for the same conditions as in (F). **H.** Average insulation score around TAD boundaries for the same conditions as in (F). **I.** Number of corner peaks identified by HICCUPS for the same conditions as in (F). **J.** *Left:* Flow cytometry profiles of CTCF-AID cells with or without auxin treatment, synchronized in G1 phase using a double thymidine block. *Right:* Chromatin fractionation and immunoblot analysis of SCC1-mEGFP and CTCF-AID cells with or without auxin treatment. Whole-cell extract (W), supernatant (S) and chromatin pellet fractions (P) were analyzed by immunoblotting using antibodies against the proteins indicated on the right. The asterisk indicates a non-specific band detected by the antibody recognizing CTCF. α-Tubulin and H3 were used as loading controls. **K.** Intra-chromosomal contact frequency distribution as a function of genomic distance for SCC1-eGFP and CTCF AID cells with or without auxin treatment. **L.** Average insulation score around TAD boundaries for the same conditions as in (G). **M.** *Top:* Contact frequency, P(s), as a function of genomic distance, s, for the same conditions as in (D). *Bottom:* Log derivatives of P(s). Arrows and dotted lines indicate peaks in the log derivatives and their positions. These correspond to the loop extrusion shoulder, which indicates the mean size of an extruded loop(Gassler et al., 2017; Polovnikov et al., 2023). Loop sizes are shown in kb. See Methods. **N.** Top and bottom: Coverage-corrected Hi-C contact matrices for the 100–132 Mb (top) and for the 25-50 Mb (bottom) regions of chromosome 2, visualized using Juicebox, in control and WAPL/PDS5A/B (WAB) depleted cells. ChIPseq tracks for the active histone mark H3K4me1 and the inactive mark H3K9me3 are shown above the Hi-C matrices. The dashed lines indicate boundaries between active and inactive regions, which correlate with the borders of the checkerboard pattern.

**Figure S2.**
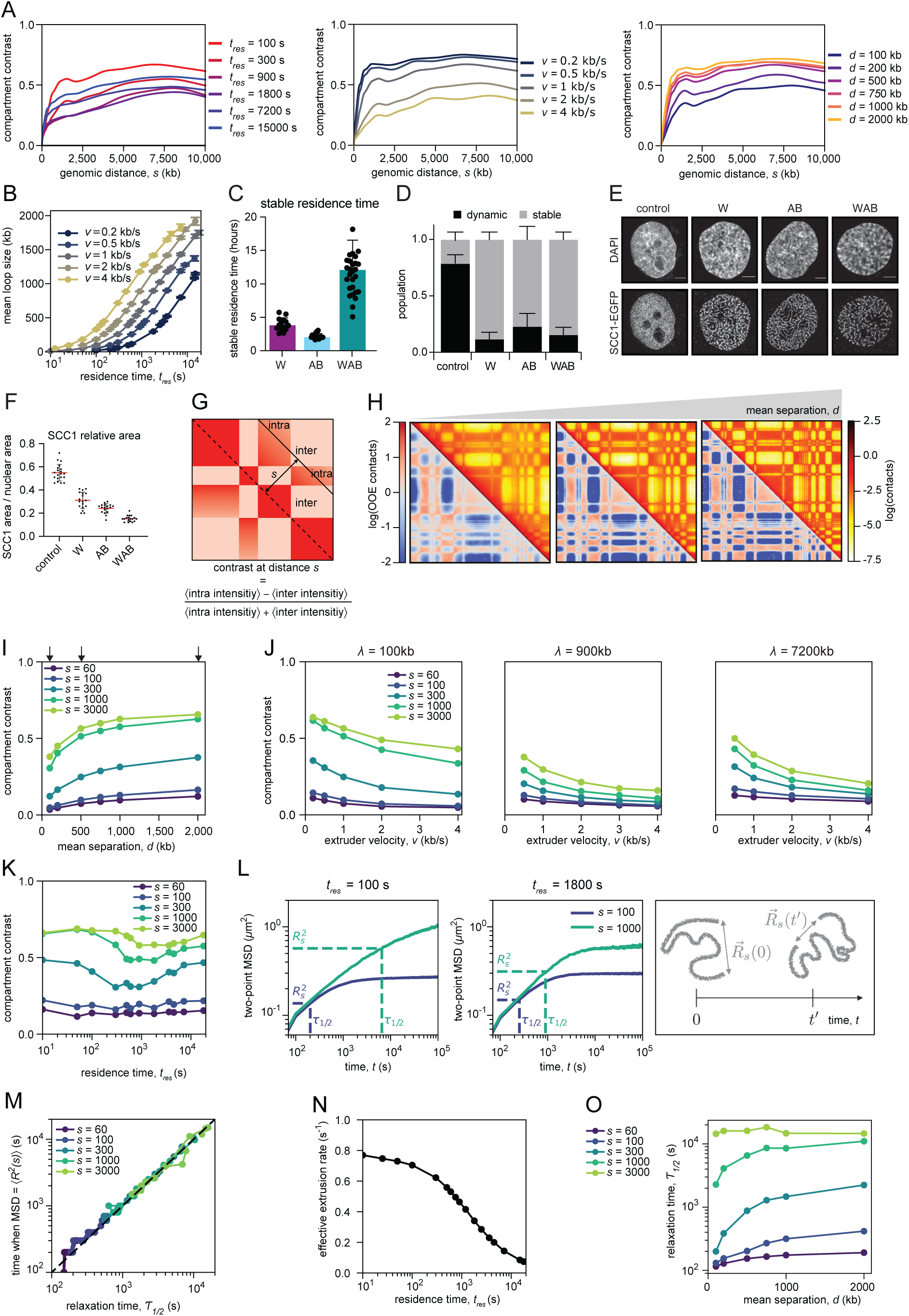
Compartments are suppressed by dynamic loop extrusion but not by the presence of static cohesin loops, related to Figure 2. **A.** *Left:* Compartment contrast versus genomic distance, *s*, from simulations for different residence times (*t_res_*). *Middle:* Compartment contrast versus genomic distance, *s*, from simulations for different extrusion velocities *(v*). *Bottom:* Compartment contrast versus genomic distance, *s*, from simulations for different mean separations between extruders (*d*). **B.** Mean extruded loop sizes as function of residence time (*t_res_*) for different extrusion velocities (*v*). **C.** Quantification of the stable residence time of SCC1-eGFP, determined by iFRAP (Figure 2C), following depletion of WAPL (W), PDS5A/B (AB) and WAPL/PDS5A/B (WAB; mean ± SD). **D.** Percentage of stable and dynamic SCC1-eGFP populations determined by iFRAP (Figure 2C) in the control cells and after depletion WAPL (W), PDS5A/B (AB) and WAPL/PDS5A/B (WAB). **E.** Airyscan images of immunofluorescence staining for DAPI and SCC1-eGFP in control, WAPL-depleted (W), PDS5A/PDS5B-depleted (AB) and WAPL/PDS5A/B- depleted cells (WAB). **F.** Vermicelli strength quantification for the conditions shown in (E), determined by dividing SCC1-eGFP signal area by the nuclear area defined by DAPI staining. **G.** Illustration of a contact map showing the computation of compartment contrast. At each diagonal distance s from the main diagonal, compartment contrast is calculated as the difference between the aggregate intra-compartment contact and inter-compartment contact frequencies (indicated by color intensity), scaled by the sum of these two values. See Methods for more details. **H.** Contact maps from simulations with varying mean separations (*d*) between loop extruders, with *d* increasing from left to right. **I.** Compartment contrast plotted against mean separation (*d*) for different genomic distances (*s*, in kb, indicated by different colors). Arrows mark the mean separations used in the simulations shown in (H). **J.** Compartment contrast plotted against extrusion velocity (*v*), with processivity, *λ=2vt_res_*, held constant. That is, residence time decreases with increasing velocity to maintain constant processivity. **K.** Compartment contrast as a function of residence time (*t_res_*) for simulations with frozen loops for different genomic distances, s. Here, residence time denotes the tres implemented in the initial active extrusion simulation performed to generate the static loop architecture to be used in the 3D polymer simulation with frozen loops (see Methods). **L.** *Left:* Two-point mean-squared displacement (MSD) for different polymer segment sizes (*s*) in simulations with two different residence times (*t_res_*). Vertical dashed lines indicate relaxation time (τ_1/2_) for each segment size. Horizontal dashed lines represent the mean-squared segment end-to-end distance, 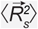. Right: Illustrations showing how the end-to-end distance of an individual polymer segment changes over time. **M.** Mean time at which the two-point MSD for a segment of size s exceeds 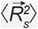 plotted against the relaxation time (t1/2) for the corresponding segment. Data are from simulations with *v* = 1 kb/s and all include all simulated tres. **N.** Effective extrusion rate, computed as the number of extruder steps per unit time per extruder in simulations, plotted against residence time (*t_res_*). **O.** Relaxation time (τ_1/2_) plotted against the mean separation (*d*) between loop extruders.

**Figure S3.**
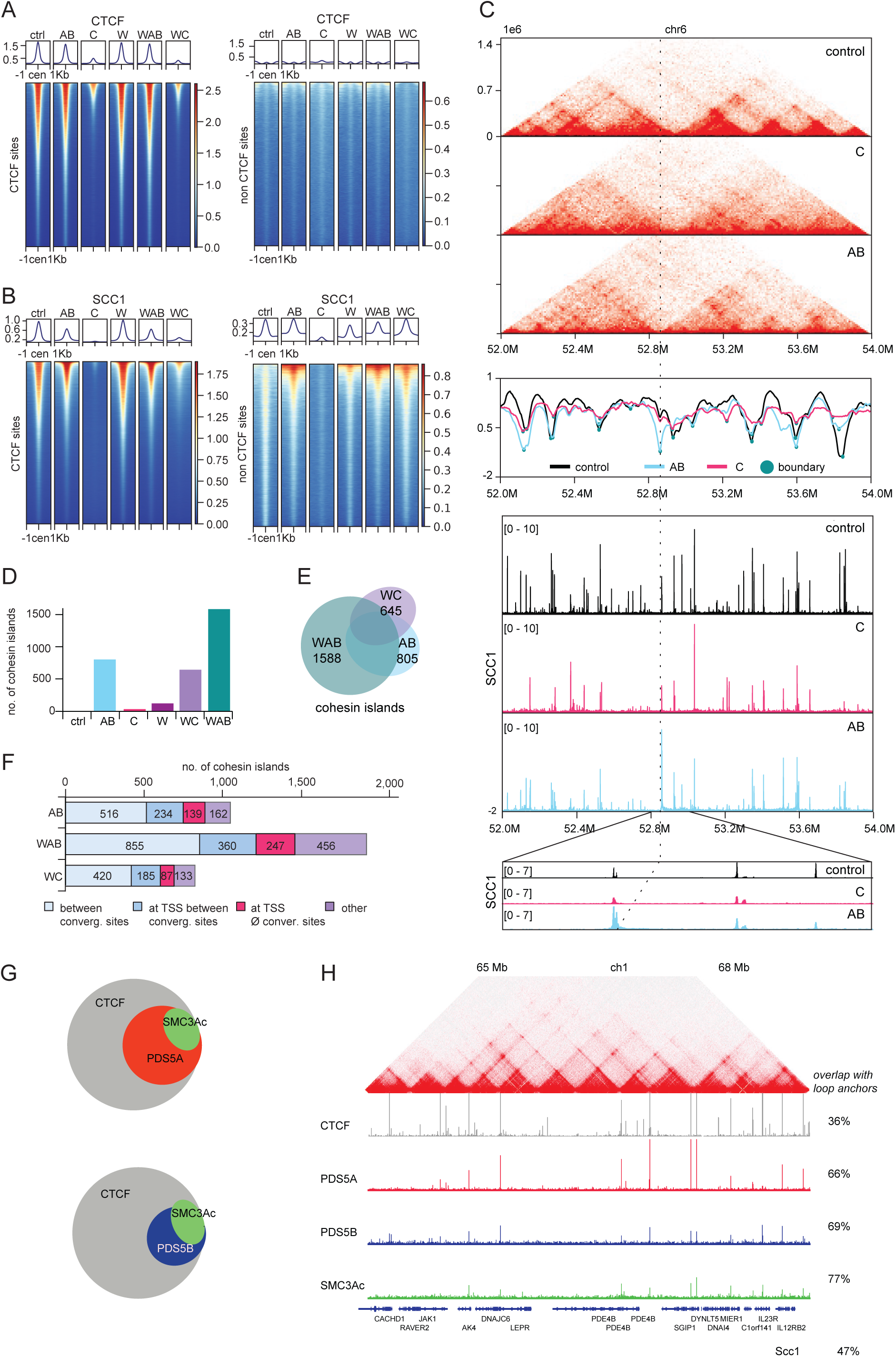
PDS5 proteins control the genomic positioning of cohesin at TAD boundaries but are dispensable for the accumulation of CTCF at its binding sites, related to Figure 3. **A.** Heatmaps of CTCF ChIP-seq signal intensities in control, PDS5A/B (AB), CTCF (C), WAPL (W), WAPL/PDS5A/B (WAB) and WAPL/CTCF (WC)-depleted cells centered on detected peaks (+/- 1 kb). CTCF coordinates represent a comprehensive compilation of all detectable peaks. Non-CTCF sites refer to SCC1 peaks (see B) at positions where CTCF is not significantly detectable. Heatmap columns are sorted by overall intensity across all conditions, line graphs above the heatmaps show the average signal. **B.** Heatmaps of SCC1 ChIP-seq signal intensities under the same depletion conditions as in (A) around the centers of detected peaks (+/- 1 kb). Cohesin sites are condition specific. Non-CTCF sites refer to SCC1 peaks at positions without significant CTCF signal. Heatmap columns are sorted by overall intensity across all conditions, line graphs above the heatmaps show the average signal. **C.** *Top:* Coverage-corrected Hi-C contact matrices for the 52-54 Mb region of chromosome 6 in control, CTCF (C) and PDS5A/B (AB) depleted cells. *Middle:* Insulation scores for the same genomic region and conditions as above. Strong boundaries were defined by strength values greater than 0.6 (turquoise circles). *Bottom:* ChIP-seq profiles of SCC1 and CTCF for the same genomic region and conditions as above. The dashed line marks a new boundary in the AB condition corresponding to a cohesin island. **D.** Number of called cohesin islands in the same depletion conditions as in (A). **E.** Venn diagram showing the overlap of cohesin island regions in WAB, WC, and AB depleted cells. **F.** Number of cohesin islands located between convergent genes, at transcription start sites (TSS) or elsewhere in the genome for the same conditions as in (A). **G.** Venn diagrams showing the overlap of CTCF, acetylated SMC3 (SMC3ac) and PDS5A or PDS5B ChIP-seq sites. **H.** Coverage-corrected Hi-C contact matrix of the 65-68 Mb region of chromosome 1 in HeLa cells. ChIP-seq signals for CTCF, eGFP- PDS5A, eGFP-PDS5B and SMC3ac are shown below the Hi-C matrix. The SMC3ac ChIP-seq data were previously published in Wutz et al., 2020. The percentage of genome-wide overlap between loop anchors identified from the Hi-C map of control HeLa cells (Wutz et al., 2020; Figure 4) and ChIP-seq binding sites are indicated on the right.

**Figure S4.**
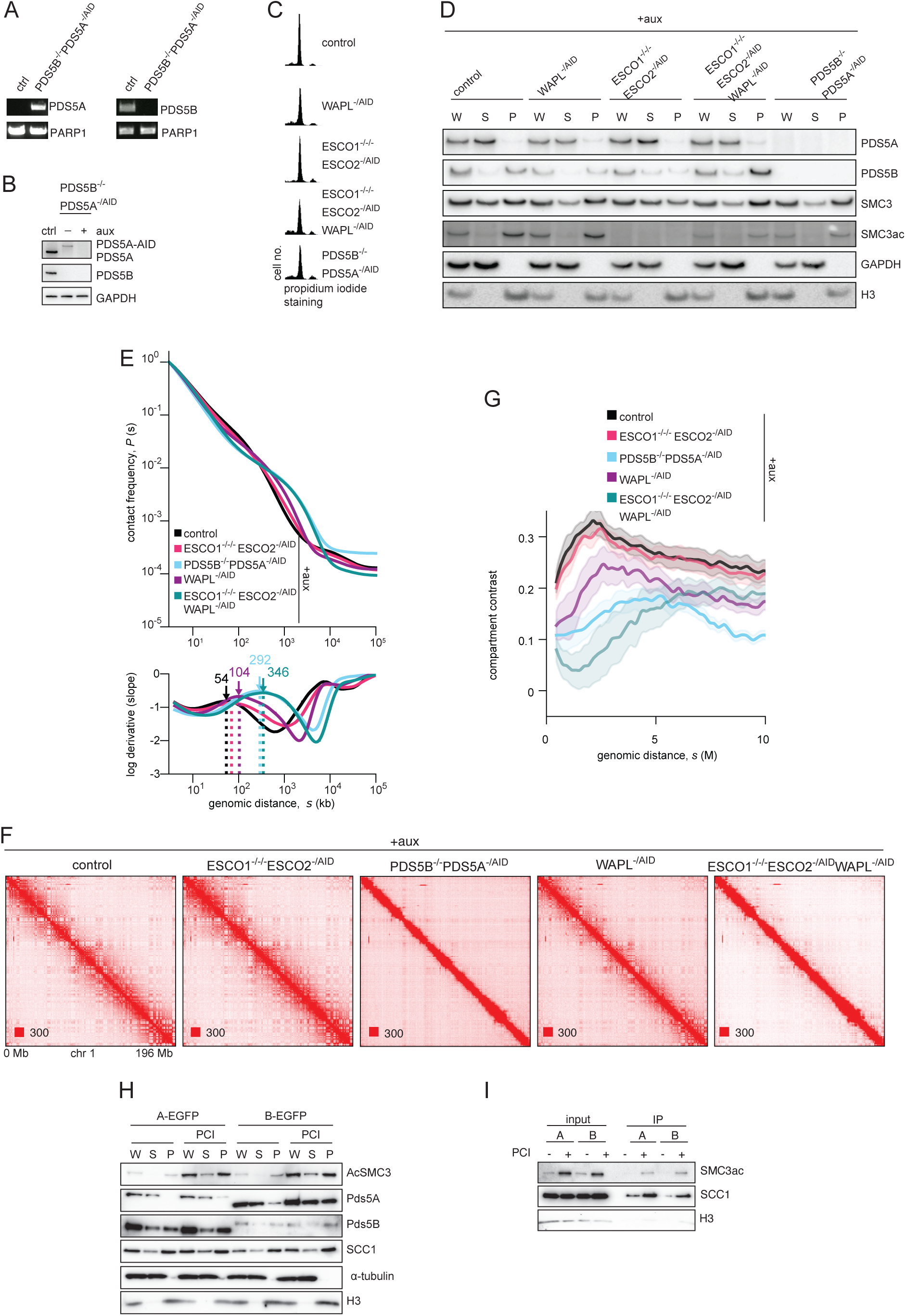
The role of PDS5 proteins in enabling cohesin acetylation is not sufficient to explain their role in CTCF boundaries, related to Figure 4. **A.** Genomic PCR genotype analysis of control and PDS5B^-/-^PDS5A^-/AID^ DT40 cell lines. The PDS5A PCR was performed using primers specific for the AID tag insertion. The PDS5B PCR was performed using PDS5B-specific primers. Primers specific to the PARP1 gene were used as a loading control. **B.** Immunoblotting analysis of whole-cell extracts from control and PDS5B^-/-^PDS5A^-/AID^ DT40 cells with or without auxin treatment. GAPDH was used as a loading control. **C.** Flow-cytometry profiles of control, WAPL, ESCO1/2, WAPL/ESCO1/2 and PDS5A/B depleted cells synchronized in late G1/early S. **D.** Chromatin fractionation and immunoblot analysis of auxin-treated WAPL^-/AID^, ESCO1^-/-/-^ESCO2^-/AID^, ESCO1^-/-/-^ES- CO^2-/AID/^/WAPL^-/AID^ and PDS5B^-/-^PDS5A^-/AID^ cells. Whole-cell extract (W), supernatant (S) and chromatin pellet fractions (P) were analyzed by immunoblotting using the antibodies against the proteins indicated on the right. GAPDH and H3 were used as loading controls. **E.** *Top:* Contact frequency, P(s), as a function of genomic distance s, for the same conditions as in (D). *Bottom:* Logarithmic derivatives of P(s). Arrows and dotted lines indicate peaks in the log derivatives and their positions, corresponding to the loop extrusion shoulder, which reflects the mean size of extruded loops(Gassler et al., 2017; Polovnikov et al., 2023). Loop sizes are shown in kb. See Methods. **F.** Coverage-corrected Hi-C contact matrices for the entire chromosome 1 visualized using Juicebox for wild type, ESCO1/ESCO2, PDS5A/PDS5B, WAPL and triple ESCO1/ESCO2/WAPL-depleted cells. **G.** Compartment contrast as a function of genomic distance (s) for control and perturbation conditions (see Fig. S2G and Methods).Shaded areas indicate standard error of the mean, calculated from the distance-dependent compartment contrasts of each individual chromosome. **H.** Chromatin fractionation and immunoblot analysis of eGFP-PDS5A and eGFP-PDS5B HeLa cells treated with the HDAC8 deacetylase inhibitor (PCI). Whole-cell extract (W), supernatant (S) and chromatin pellet fractions (P) were analyzed by immunoblotting using the antibodies against the proteins indicated on the right. **I.** Immunoblot analysis of immunoprecipitates from eGFP-PDS5A and eGFP-PDS5B cells treated with PCI. Immunoprecipitation was performed using an anti-GFP antibody, and immunoblotting was carried out with the antibodies against the proteins indicated on the right.

**Figure S5.**
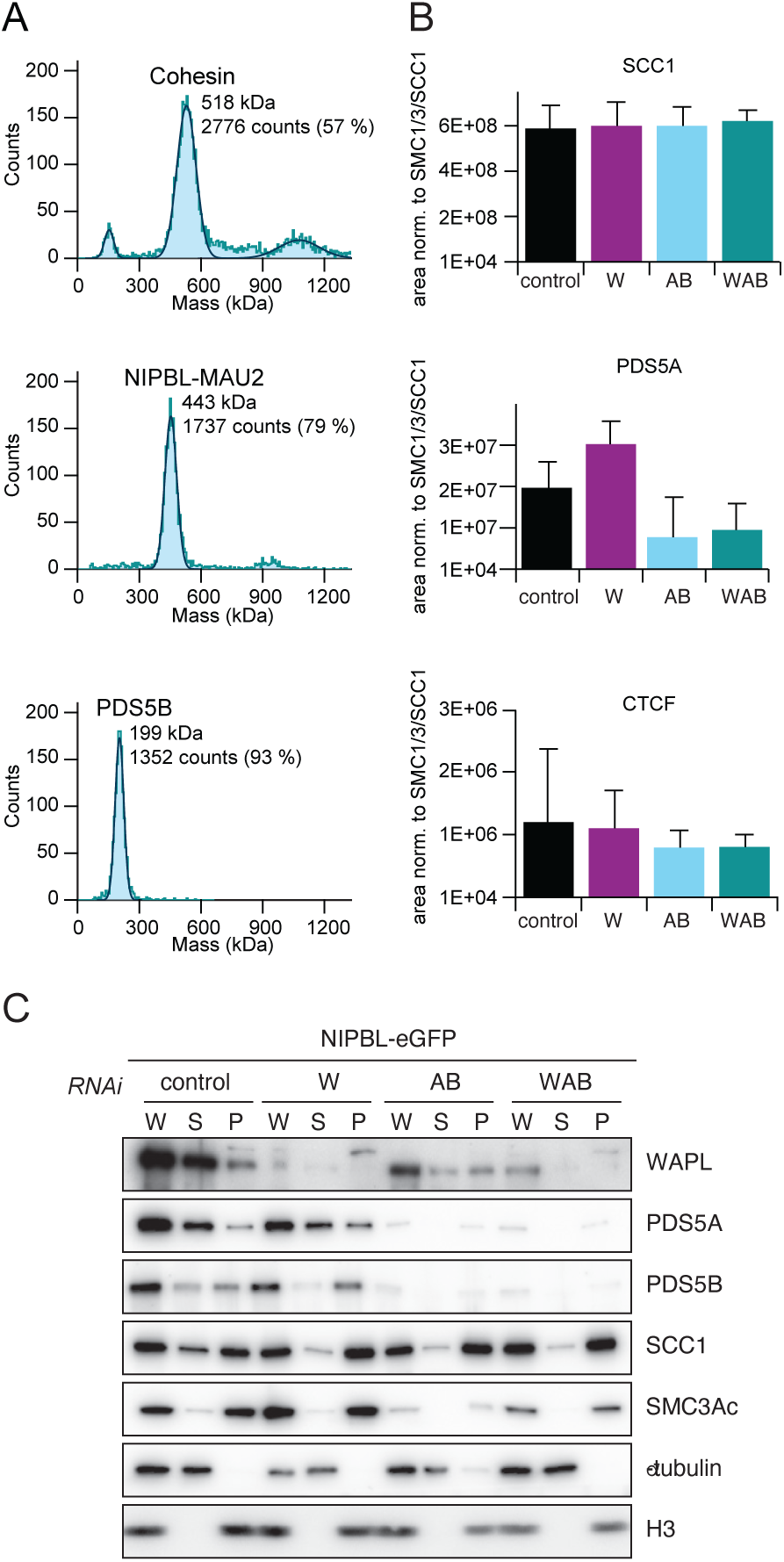
PDS5 proteins limit the amount of NIPBL bound to cohesin and the residence time of NIPBL on chromatin, related to Figure 5. **A.** Representative mass histograms (blue) and Gaussian fits of cohesin, NIPBL-MAU2 and PDS5B. Observed molecular masses and the percentage of total counts corresponding to the indicated peak are shown. **B.** Amount of PDS5A and CTCF bound to cohesin was determined by quantitative mass spectrometry from material immunoprecipitated using anti-GFP antibodies from SCC1- GFP cells synchronized in G1 and depleted for WAPL (W), PDS5A/B (AB) and WAPL/PDS5A/B (WAB). Peptide coverage was calculated relative to the total number of SMC3, SMC1 and SCC1 peptides. **C.** Chromatin fractionation and immunoblot analysis of NIPBL-eGFP HeLa cells treated with WAPL (W) and PDS5A/B (AB) and WAPL/PDSS5A/B (WAB) siRNAs. Following RNAi, whole-cell extract (W), supernatant (S) and chromatin pellet fractions (P) were analyzed by immunoblotting, using the antibodies against the proteins indicated on the right.

**Figure S6.**
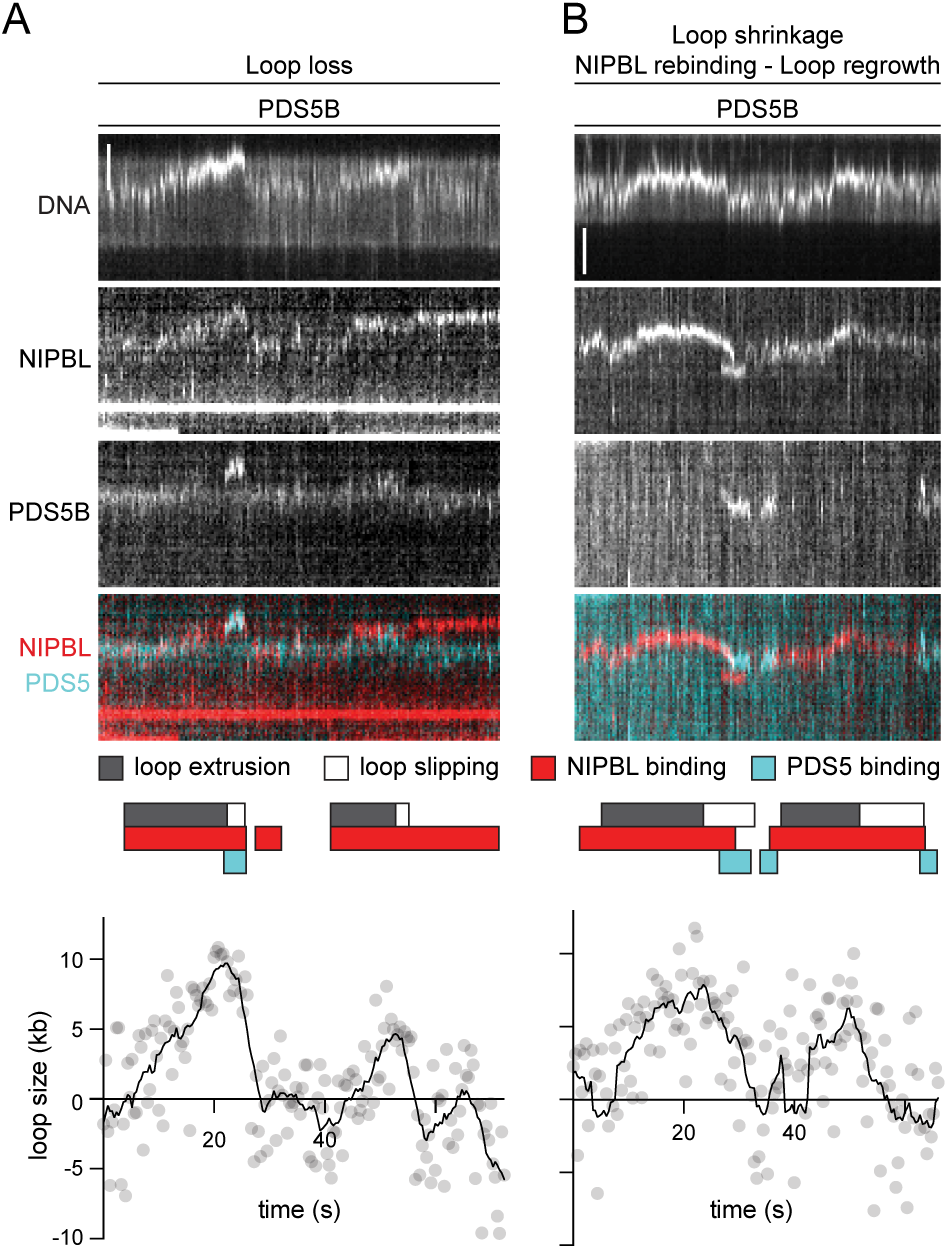
Additional examples of NIPBL-ΔN / PDS5B exchange and loop loss following NIP- BL-ΔN and cohesin dissociation, related to Figure 6. **A.** Example kymographs of NIPBL-ΔN/PDS5B exchange during cohesin-mediated loop extrusion. PDS5B binds at the loop base at 23.2 s. NIPBL-ΔN dissociates and the loop becomes undetectable at 27.2 s. NIPBL-ΔN rebinds at 28.4 s. Loop size graph dots represent raw data, solid lines represent a smoothed version using a Savitzky-Golay filter with 5 neighbors and order 0. Phases of loop extrusion, loop slipping, NIPBL binding and PDS5 binding are represented by colored blocks above the loop size graph. DNA was stained with SYTOX Green. NIPBL-ΔN was labeled with ATTO550; PDS5B was labeled with JF646. Scale bar, 2 µm. **B.** As (A) except in this example, NIPBL-ΔN is first displaced by PDS5B and then rebinds to the loop base and displaces PDS5B.

**Figure S7.**
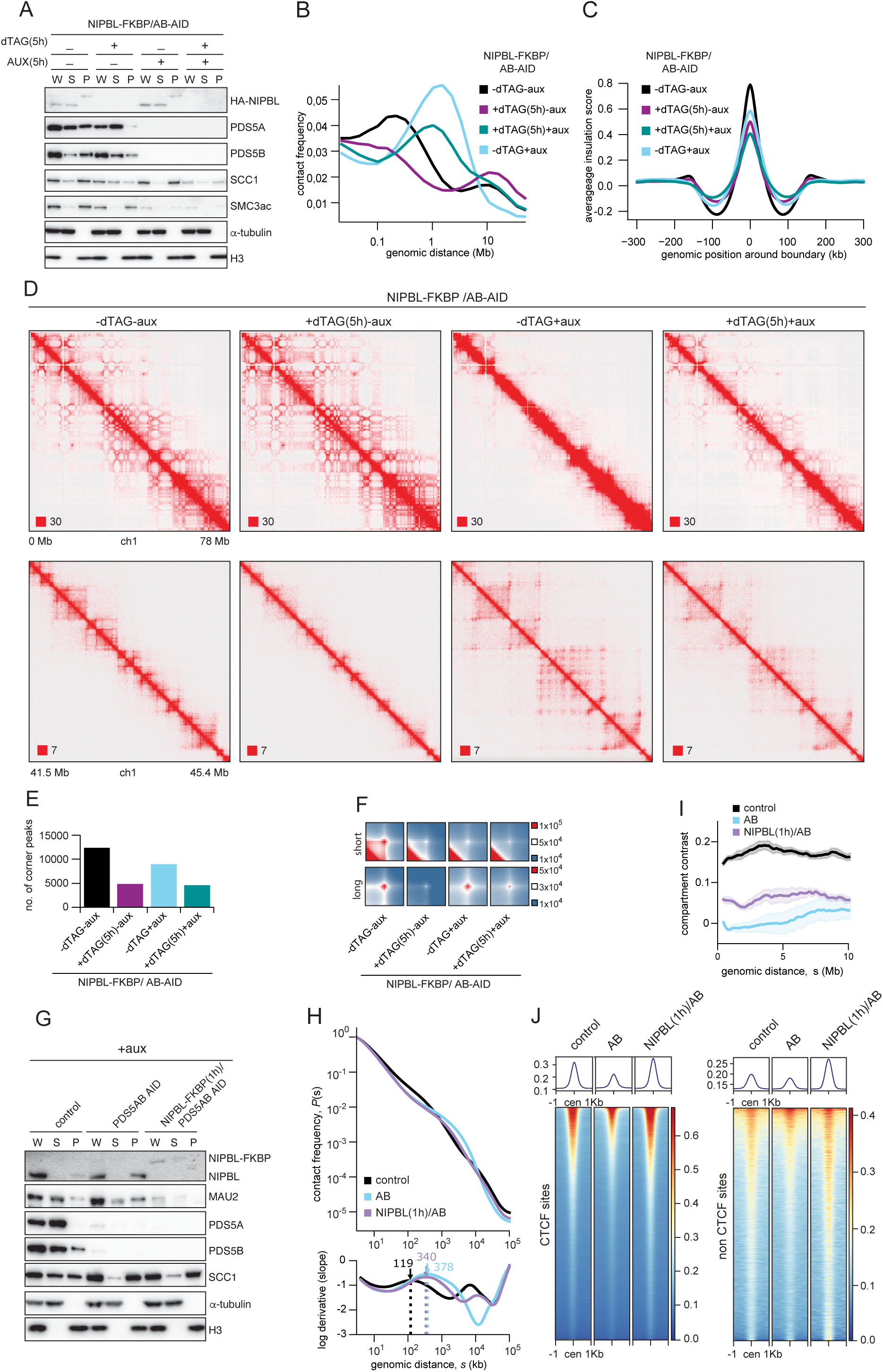
PDS5 proteins strengthen CTCF boundaries by limiting the lifetime of cohesin-NIPBL complexes, related to Figure 7. **A.** Chromatin fractionation and immunoblot analysis of NIPBL-FKBP/PDS5AB (AB)-AID cells treated with auxin and/or dTAG for 5h. Whole-cell extract (W), supernatant (S) and chromatin pellet fractions (P) were analyzed by immunoblotting using antibodies against the proteins indicated on the right. a-tubulin and H3 were used as loading controls. **B.** Intra-chromosomal contact frequency distribution as a function of genomic distance for the same conditions as in (A). **C.** Average insulation scores around TAD boundaries for the same conditions as in (A). **D.** Coverage-corrected Hi-C contact matrices for the 0-78Mb (top) and 41.5-45.4 Mb (bottom) regions of chromosome 1. Matrices were plotted using Juicebox for the same conditions as in (A). **E.** Number of corner peaks identified by HICCUPS for the same conditions as in (A). **F.** Total contact counts around corner peaks are shown for different conditions. Short corner peaks are those ≤350 kb identified in control cells. Long corner peaks are those >350 kb identified in WAPL-depleted cells. Matrices are centered at the corresponding loop anchor sites in vertical and horizontal orientation. **G.** Chromatin fractionation and immunoblot analysis of NIPBL-FKBP/PDS5AB-AID Hela cells treated with dTAG for 1hour compared to control and AB depleted cells (auxin treatment was applied for 5h in all cases). Whole-cell extract (W), supernatant (S) and chromatin pellet fractions (P) were analyzed by immunoblotting using antibodies against the proteins indicated on the right. a-tubulin and H3 were used as loading controls. **H.** *Top:* Contact frequency, P(s), as a function of genomic distance, s, for the indicated conditions. *Bottom:* Logarithmic derivatives of P(s). Arrows and dotted lines mark the peaks in the derivatives, which correspond to the loop extrusion shoulder, indicating the mean size of extruded loops. Loop sizes are shown in kb. See Methods. **I.** Compartment contrast as a function of genomic distance, s (see Fig. S1M and Methods), for control and perurbed conditions. The shaded area reprisents standard error of the mean, calculated from the mean compartment contrasts of each individual chromosome. **J.** Heatmaps of SCC1 ChIP-seq signal intensities in the same depletion situations as in (G), centered on detected peaks +/- 1 kb. Cohesin sites are sample-specific. “Non-CTCF sites” refer to SCC1 peaks at locations significant CTCF signal. Heatmap columns are sorted by average signal intensity across all samples, line graphs above each heatmap show the average profile.

**Table S1.**
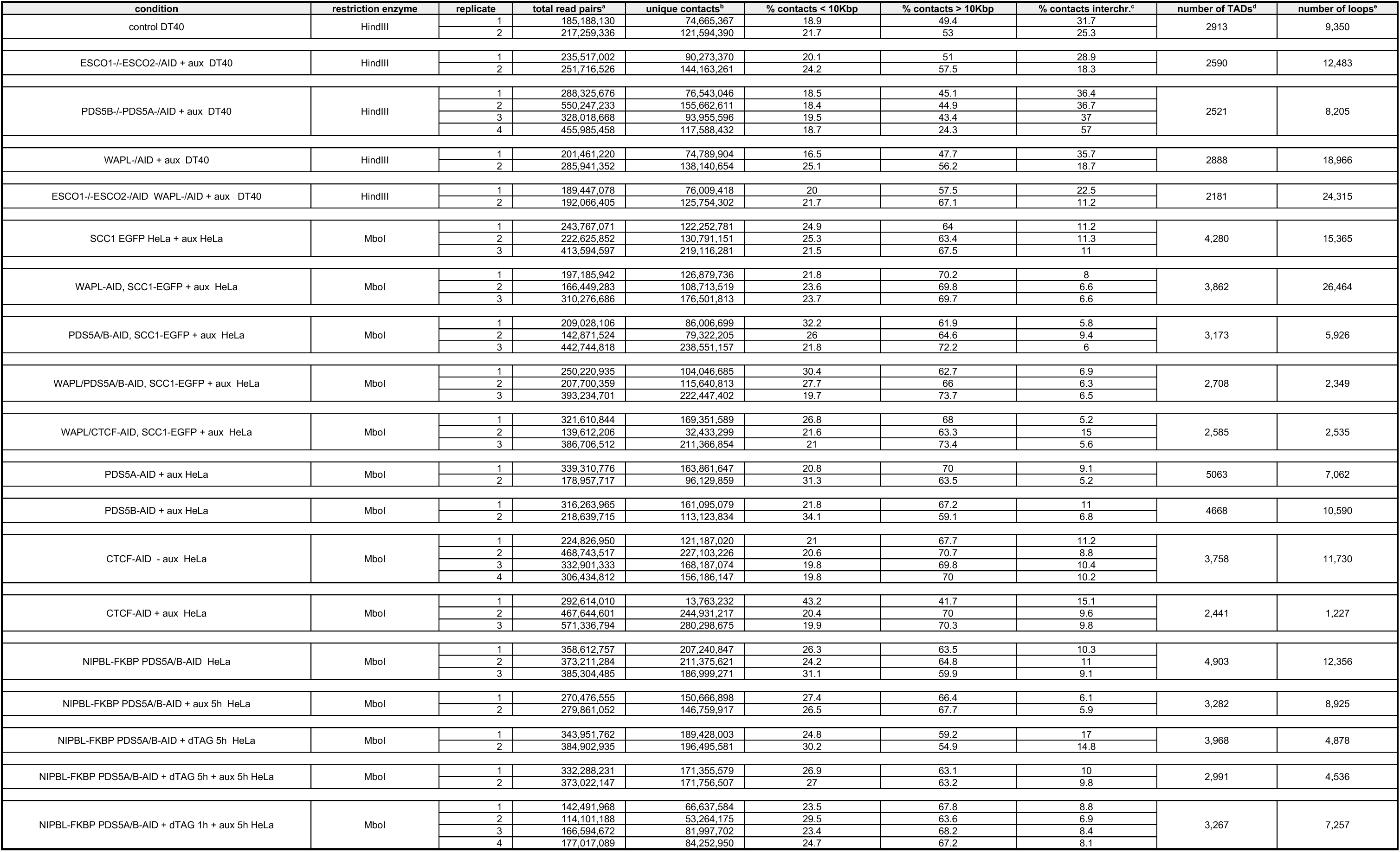
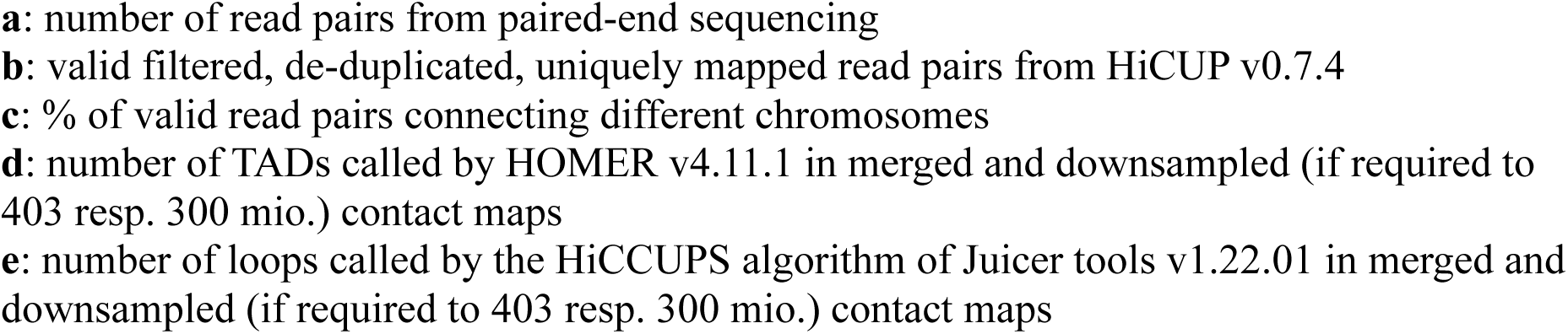
Summary statistics for Hi-C data sets generated in this study, related to Figure 1 and 7.

**Table S2.**
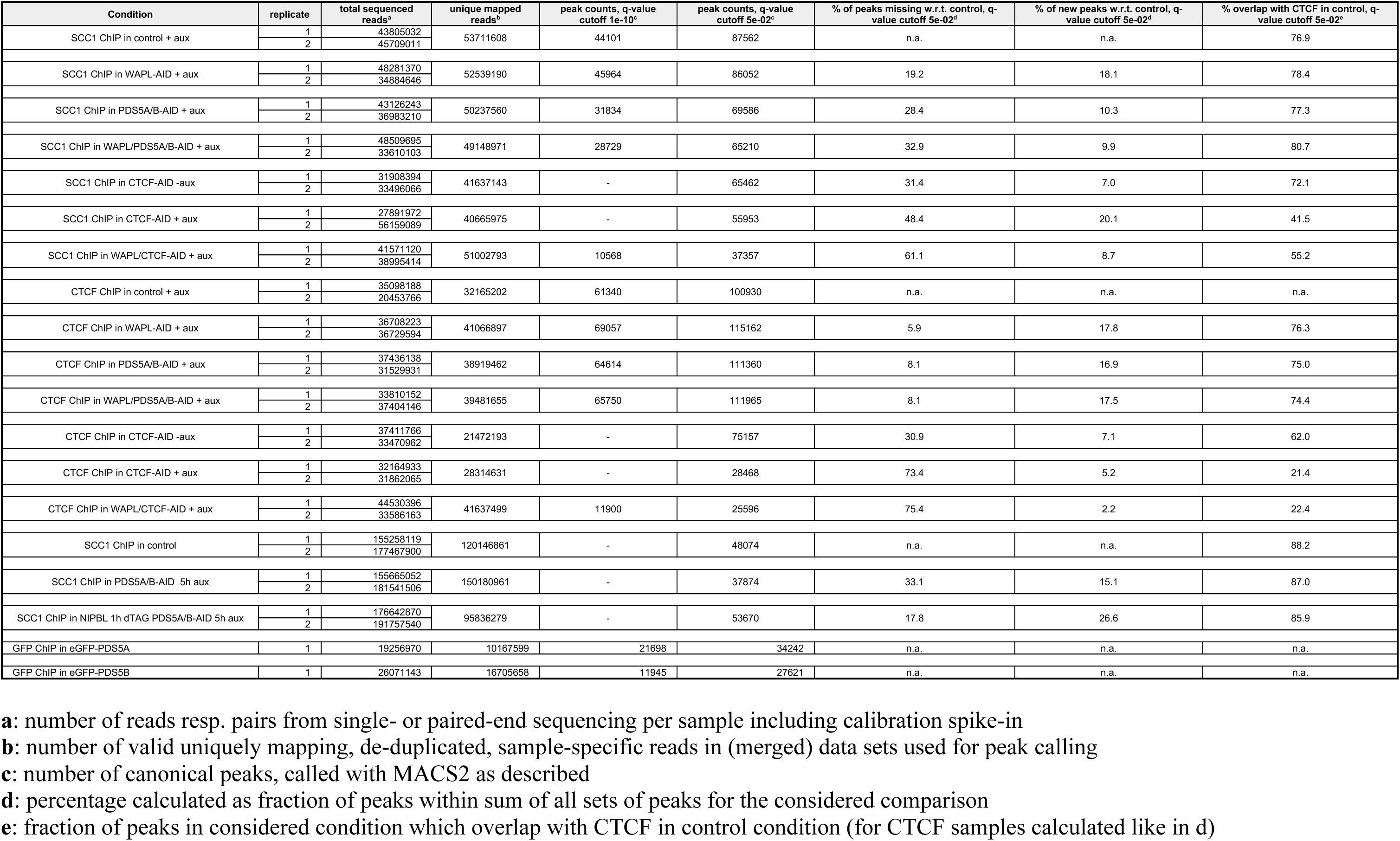
Summary statistics for ChIP-Seq data sets generated in this study, related to Figure 3 and 7.

